# Effects of lysine deacetylation inhibition alone or in combination with arimoclomol on TDP-43 proteinopathy

**DOI:** 10.1101/2024.12.15.628528

**Authors:** Serena Scozzari, Stefano Fabrizio Columbro, Monica Favagrossa, Massimo Tortarolo, Alfredo Cagnotto, Mario Salmona, Giovanni De Marco, Caterina Bendotti, Andrea Calvo, Laura Pasetto, Valentina Bonetto

**Author notes:** These are co-corresponding authors email: Laura Pasetto or Valentina Bonetto. These are co-first authors (Serena Scozzari, Stefano Fabrizio Columbro).

## Abstract

Cytoplasmic inclusions containing TAR DNA-binding protein 43 kDa (TDP-43) are recognized as a major pathological feature of amyotrophic lateral sclerosis (ALS) and frontotemporal dementia. Peptidyl-prolyl cis-trans isomerase A (PPIA) interacts with TDP-43 and influences its aggregation and function. This interaction is facilitated by PPIA lysine acetylation. Here, we investigated whether restoring lysine acetylation homeostasis exerts protective effects on TDP-43 proteinopathy *in vitro* and *in vivo*and how this relates with PPIA. We found that vorinostat/SAHA, a broad-spectrum histone deacetylase (HDAC) inhibitor that increases PPIA acetylation, is able to reverse TDP-43 mislocalization in a cellular model of TDP-43 proteinopathy. We confirmed its effects in peripheral blood mononuclear cells from ALS patients and explored its impact on TDP-43 proteinopathy and PPIA acetylation in the Thy1-hTDP-43 mouse model. Thy1-hTDP-43 mice treated with SAHA showed a delayed onset of TDP-43 pathology, associated with PPIA nucleus-cytoplasm redistribution, lower neurodegeneration and neuroinflammation, and improved neuromuscular function markers. However, these effects were transient. When combined with arimoclomol, a heat shock protein co-inducer, a mitigation of the neurodegeneration was sustained. A synergistic effect was observed in periphery, greatly enhancing tubulin acetylation and reducing phosphorylated TDP-43 accumulation in the sciatic nerve and acetylcholine receptor γ-subunit expression in gastrocnemius muscle. This study suggests that HDAC inhibition could be beneficial in restoring TDP-43 localization and function through multiple mechanisms, including modulation of PPIA acetylation. The combination of HDAC inhibition and arimoclomol shows a synergistic effect *in vivo* and has potential as a therapeutic approach for patients.

## 1 Introduction

Cytoplasmic inclusions containing TAR DNA-binding protein 43 kDa (TDP-43) were originally discovered in affected central nervous system (CNS) regions of patients with amyotrophic lateral sclerosis (ALS) and frontotemporal lobar degeneration (Neumann *et al*. 2006). Since then, several studies have contributed to establish TDP-43 inclusions as a major pathological component of ALS and frontotemporal dementia (FTD), and mutations in *TARDBP*, the gene encoding for TDP-43, have been found to cause familial ALS, ALS-FTD and FTD, although rarely. More recently, intraneuronal cytoplasmic inclusions of TDP-43 have also been seen in the brain of more than 50% of Alzheimer’s (AD) disease patients (McAleese *et al*. 2017), about 25% of elderly individuals, as limbic-predominant age-related TDP-43 encephalopathy neuropathological changes (Nelson *et al*. 2019), and 40% of subjects with chronic traumatic encephalopathy linked to repetitive head injuries (Nicks *et al*. 2023), Parkinson’s (Yamashita *et al*. 2022) and Huntington’s disease (Sanchez *et al*. 2021). These later discoveries have broadened the range of TDP-43-related conditions (de Boer *et al*. 2020) and underscored the importance of understanding the molecular mechanisms behind this pathology for developing effective treatments.

TDP-43 is an RNA-binding protein normally localized in the nucleus, also capable of shuttling to the cytoplasm, to govern various cellular functions such as transcriptional repression, pre-mRNA splicing, mRNA stability, microRNA biogenesis, RNA transport, and translational regulation (Ratti and Buratti 2016). While the precise pathological function of TDP-43 remains elusive, mounting evidence suggests that its depletion from the nucleus and its accumulation in the cytoplasm can result in both gain and loss of functions. Due to its aggregation propensity, cells tightly regulate TDP-43 levels by an auto-regulatory feedback loop involving its own mRNA transcript (Ayala *et al*. 2011; Polymenidou *et al*. 2011) and, upon stress conditions, activate protective mechanisms by which transiently store TDP-43 into liquid-liquid phase-separated droplets in the nucleus and the cytoplasm (Yu *et al*. 2021; Lu *et al*. 2022; Wang *et al*. 2020). Under prolonged stress conditions, the coping response fails and TDP-43 accumulates as aggregates mainly in the cytoplasm likely causing nucleocytoplasmic transport impairment, clearance of nuclear TDP-43, impaired splicing function, deficient axonal transport and cell death (Gasset-Rosa *et al*. 2019; Fazal *et al*. 2021; Chou *et al*. 2018; Fratta *et al*. 2018).

Peptidyl-prolyl cis-trans isomerase A (PPIA), also known as cyclophilin A, a foldase and molecular chaperone, interacts with TDP-43 and impacts on its aggregation (Lauranzano *et al*. 2015; Pasetto *et al*. 2021). Under stress conditions, PPIA has been found associated with membrane-less assemblies in the cytoplasm (Curdy *et al*. 2023; Lu *et al*. 2022; Xiang *et al*. 2015) and in the nucleus (Yu *et al*. 2021), and there is evidence that it regulates liquid-liquid phase separation of its substrates (Babu *et al*. 2022; Maneix *et al*. 2024). Moreover, it has been found sequestered in protein aggregates in ALS patients and mouse models (Lauranzano *et al*. 2015; Basso *et al*. 2009). PPIA knockout mice develop a neurodegenerative disease which resembles ALS-FTD with marked TDP-43 pathology (Pasetto *et al*. 2021). PPIA depletion in SOD1^G93A^ mice induces TDP-43 aggregation and accelerates disease progression (Lauranzano *et al*. 2015). The interaction between PPIA and TDP-43 occurs at the Gly-Pro dipeptide motif (residues 348-349) in the C-terminal Gly-rich domain of TDP-43 that has prion-like properties, and is favored by PPIA acetylation at K125 (Lauranzano *et al*. 2015). PPIA Lys-acetylation is low in cells under stress conditions and in peripheral blood mononuclear cells (PBMCs) of ALS patients (Lauranzano *et al*. 2015), where signs of TDP-43 proteinopathy have been found (De Marco *et al*. 2011; Luotti *et al*. 2020). Moreover, we identified a PPIA loss-of-function K76E mutation, at a Lys susceptible to acetylation (Choudhary *et al*. 2009), in a patient with sporadic ALS (Pasetto *et al*. 2021). However, it is not known if modulating PPIA acetylation has an effect on TDP-43 aggregation and function.

The level of Lys-acetylation decreases globally in neurons during neurodegeneration, reflecting an impairment in the acetylation homeostasis (Saha and Pahan 2006). The removal of acetyl groups is mediated by histone deacetylases (HDACs), a superfamily of enzymes grouped in four classes, now also named lysine deacetylases in consideration of the numerous non-histone protein targets recently identified (Narita *et al*. 2019). HDAC inhibitors (HDACis) have shown to have interesting neuroprotective effects for ALS and in general for neurodegenerative diseases (Shukla and Tekwani 2020; Klingl *et al*. 2021). Nevertheless, first generation HDACis tested in clinical studies for ALS, valproic acid and sodium phenylbutyrate, had no beneficial effects (Piepers *et al*. 2009; Cudkowicz *et al*. 2009), probably because having a broad spectrum of action have also harmful effects (Boutillier *et al*. 2019; Pigna *et al*. 2019). Novel, more specific, molecules are now available. An HDAC3 inhibitor regulated the expression of the amyloid precursor protein and promoted neuroprotective pathways in an AD mouse model (Davis *et al*. 2024). HDAC6 inhibitors reversed axonal transport defects in FUS mice and in motor neurons derived from ALS patients, carrying FUS and TDP-43 mutations (Fazal *et al*. 2021; Guo *et al*. 2017; Rossaert *et al*. 2019). A SIRT1 inhibitor counteracted mutant SOD1 toxicity in neuronal cells (Valle *et al*. 2014). Interestingly, the broad-spectrum inhibitor suberoylanilide hydroxamic acid (SAHA), also known as vorinostat, and the HDAC1/3 inhibitor RGFP109 can induce heat shock protein (HSP) response in motor neurons expressing variants linked to familial ALS (Kuta *et al*. 2020; Fernández Comaduran *et al*. 2024). Moreover, a stronger and more sustained HSP response is obtained when these compounds are combined with the HSP co-inducer arimoclomol (Kuta *et al*. 2020; Fernández Comaduran *et al*. 2024).

In this study, we investigated the effect of lysine deacetylation inhibition in an *in vitro* model, in a mouse model of TDP-43 proteinopathy and in PBMCs of ALS patients. The aim was to determine how changes in PPIA acetylation correlate with TDP-43 proteinopathy and identify novel therapeutic strategies.

## 2 Methods

### 2.1 Drugs

Drugs for cells and mice treatments were purchased from MedChemExpress and were as follows: SAHA (cat. no. HY-10221), RGFP109 (cat. no. HY-16425), RGFP966 (cat. no. HY-13909); ACY-738 (cat. no. HY-19327); tubastatin A (cat. no. HY-13271A), selisistat (cat. no. HY-15452), arimoclomol maleate (cat. no. HY-106443A).

### 2.2 Antibodies

Antibodies used for immunoblot (dot/WB) and capillary electrophoresis immunoassay (CEI) on Jess (Protein Simple, Biotechne) were as follows: rabbit polyclonal anti-C-terminal TDP-43 antibody (1:2500 for immunoblot, 1:20 for CEI; Proteintech; RRID: AB_2200505), rabbit polyclonal anti-acetyl-PPIA (Lys125) antibody (1:250 for immunoblot, 1:10 for CEI; developed in-house), rabbit polyclonal anti-PPIA antibody (1:2500 for immunoblot, 1:40 for CEI; Proteintech; RRID: AB_2237516), mouse monoclonal anti-acetyl alpha-tubulin (Lys40) antibody (1:1000 for immunoblot; Sigma-Aldrich; RRID: AB_609894), mouse monoclonal anti-alpha tubulin antibody (1:1000 for immunoblot; Santa Cruz Biotechnology; RRID: AB_628412); mouse monoclonal anti-phospho-Ser409/410 TDP-43 antibody (pTDP-43) (1:2000 for immunoblot; Cosmo Bio Co., LTD; RRID: AB_1961900), rabbit polyclonal anti-HSP70/HSPA1A antibody (specific for stress-inducible and constitutive forms) (1:20000 for immunoblot; Proteintech; RRID: AB_2264230), mouse monoclonal anti-lamin A/C antibody (1:500 for immunoblot; Millipore; RRID:AB_94752), mouse monoclonal anti-GAPDH antibody (1:10000 for immunoblot; RRID: AB_2107426), mouse monoclonal anti-HDAC1 antibody (1:20 for CEI; Cell Signaling Technology; RRID: AB_10612242), rabbit monoclonal anti-HDAC3 antibody (1:20 for CEI; Cell Signaling Technology; RRID: AB_2800047), mouse monoclonal anti-Myc Tag antibody (1:3000 for immunoblot; Millipore; RRID: AB_11211891).

### 2.3 *In vitro* model of TDP-43 pathology and treatments

HEK293 cells were cultured in DMEM GlutaMAX (Dulbecco’s Modified Eagle Medium with L-Glutamine) (cat. no. 61965026, Gibco) containing 10% fetal bovine serum (FBS) (cat. no. A5670701, Gibco), 1% non-essential amino acids (cat. no. 11140-50, Gibco) and 1% penicillin/streptomycin (cat. no. 15140-122, Gibco) and maintained at 37 °C in a humidified atmosphere and 5% CO_2_. For the *in vitro* experiments performed in this study, HEK293 cells were used between passages 5 and 15 post-thaw to minimize phenotypic drift. The cell line was purchased from ATCC (CRL-1573) and was not further authenticated upon receipt. HEK293 cells are not listed as commonly misidentified by the International Cell Line Authentication Committee. To induce TDP-43 pathology, HEK293 cells were exposed to a chronic nutrient starvation protocol, essentially as previously described (Reineke *et al*. 2018). Briefly, cells were seeded in a complete medium for 24 hours, becoming 70% confluent; the following day, FBS deprived medium was replaced, and nutrient starvation was carried out for 72 hours. The cellular model was used as a biological platform for the pharmacological testing. Cells were seeded in a complete medium and subjected to overnight pre-treatment with 0.6 µM or 1 µM HDACis: SAHA, RGFP109, RGFP966, ACY-738, tubastatin A, selisistat, or vehicle. The following day, cells were exposed to the chronic nutrient starvation protocol, collected after 72 hours, and stored as pellet at −70 °C until further analysis. For each HDACi, we performed multiple independent experiments including control untreated cells (Ctrl), starved cells treated with vehicle (Veh) and starved cells treated with the HDACi; for each experiment, TDP-43 immunoreactivity of treated conditions was normalized to the respective Ctrl cells loaded within the same blot, thereby ensuring a valid comparison of HDACi effects across experiments.

### 2.4 ALS patients

Informed written consent was obtained from all participants involved and the study was approved by the ethics committee of Azienda Ospedaliero-Universitaria Città della Salute e della Scienza (Prot. n° 0062147), Turin, Italy. The diagnosis of ALS was based on a detailed medical history and physical examination, and confirmed by electrophysiological evaluation. The patients were screened for mutations in the SOD1, C9orf72, TARDBP, and FUS genes. Inclusion criteria for ALS patients were: (i) 18–85 years old; and (ii) had a diagnosis of definite, probable or laboratory-supported probable ALS, according to revised El Escorial criteria. Exclusion criteria were: (i) diabetes or severe inflammatory conditions; (ii) active malignancy; and (iii) pregnancy or breast-feeding. ALS patients underwent a battery of neuropsychological tests and were classified according to the consensus criteria for the diagnosis of frontotemporal cognitive and behavioral syndromes in ALS. We analyzed clinical samples from nine ALS patients and 18 age- and sex-matched healthy controls. The characteristics of the patients and controls are described in Table 1.

**TABLE 1.**
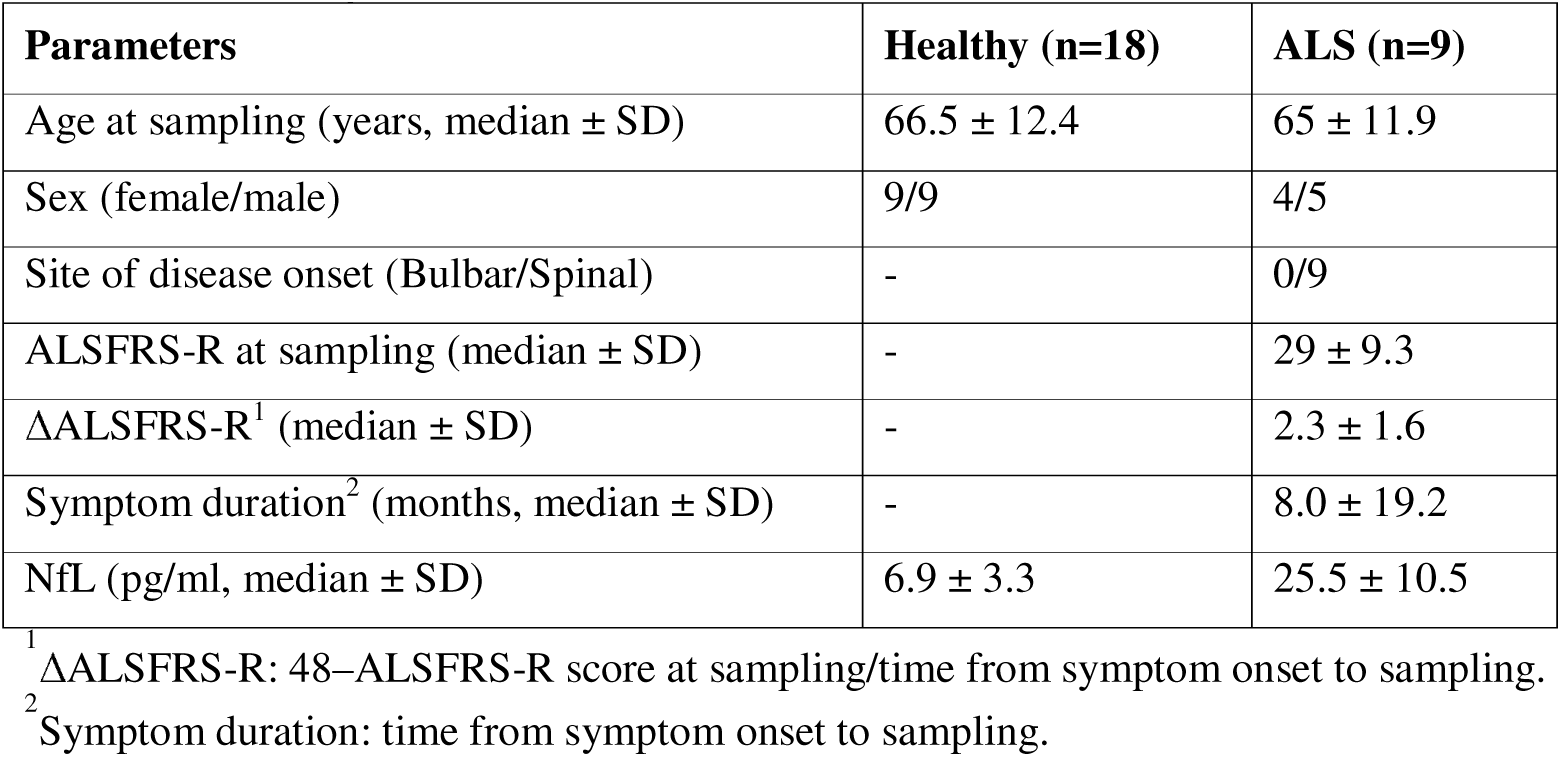
Demographic and clinical characteristics of ALS patients and controls.

| Parameters | Healthy (n=18) | ALS (n=9) |
| --- | --- | --- |
| Age at sampling (years, median ± SD) | 66.5 ± 12.4 | 65 ± 11.9 |
| Sex (female/male) | 9/9 | 4/5 |
| Site of disease onset (Bulbar/Spinal) | - | 0/9 |
| ALSFRS-R at sampling (median ± SD) | - | 29 ± 9.3 |
| ΔALSFRS-R<sup>1</sup> (median ± SD) | - | 2.3 ± 1.6 |
| Symptom duration<sup>2</sup> (months, median ± SD) | - | 8.0 ± 19.2 |
| NfL (pg/ml, median ± SD) | 6.9 ± 3.3 | 25.5 ± 10.5 |

<sup>1</sup> $\Delta$ ALSFRS-R: 48–ALSFRS-R score at sampling/time from symptom onset to sampling.
<sup>2</sup> Symptom duration: time from symptom onset to sampling.

### 2.5 Human PBMC isolation and treatment

Blood was drawn by standard venipuncture at the clinical center, collected in 4-ml BD Vacutainer® CPT™ Mononuclear Cell Preparation Tubes - Sodium Citrate (cat. no. 362781, BD) and PBMCs were isolated following the manufacturer’s protocol. PBMCs were then subjected to three additional washing steps in phosphate buffer saline and centrifugation at 300 x *g* for 15 min at RT. PBMCs were stored at −70 °C until further analysis, as pellet for biomarkers analysis or in 90% FBS and 10% dimethyl sulfoxide for *in vitro* experiments. For PBMCs culture and treatment, cells were quickly thawed in a 37 °C water bath, diluted 1:10 in warm RPMI medium 1640 (cat. no. 21870076, Gibco) and collected by centrifugation at 300 x *g* for 15 min at RT. Cells were cultured at a final concentration of 1100000 cells/ml in complete RPMI medium 1640, containing 10% FBS (cat. no. S1810-500, MicroGem), 1% L-glutamine (cat. no. FA30WX0550100, Carlo Erba Reagents) and 1% penicillin/streptomycin (cat. no. 15140-122, Gibco). PBMCs isolated from each patient were split in two cultures, one subjected to overnight treatment with 1 µM SAHA, the other group treated with vehicle; cells were incubated overnight at 37°C in a humidified atmosphere and 5% CO_2_.

### 2.6 Thy1-hTDP-43 mouse model

Procedures involving animals and their care were conducted in conformity with the following laws, regulations, and policies governing the care and use of laboratory animals: Italian Governing Law (D.lgs 26/2014; Authorization 19/2008-A issued 6 March, 2008 by Ministry of Health); Mario Negri Institutional Regulations and Policies providing internal authorization for persons conducting animal experiments (Quality Management System Certificate, UNI EN ISO 9001:2015 - Reg. N° 6121); the National Institutes of Health’s Guide for the Care and Use of Laboratory Animals (2011 edition), and European Union directives and guidelines (EEC Council Directive, 2010/63/UE). The Mario Negri Institutional Animal Care and Use Committee and the Italian Ministry of Health (Direzione Generale della Sanità Animale e dei Farmaci Veterinari, Ufficio 6) prospectively reviewed and approved the animal research protocols of this study (protocol N° 9F5F5.139 and N° 9F5F5.276, authorisation N° 688/2019-PR and N° 87/2025-PR) and ensured compliance with international and local animal welfare standards. Our Thy1-hTDP-43 mouse line is on a homogeneous C57BL/6J background and derives from the B6;SJL-Tg(Thy1-TARDBP)4Singh/J line originally obtained from The Jackson Laboratory (cat. no. 012836; Bar Harbor, ME, USA), which expresses 2 copies of human *TARDBP* under the control of a murine Thy-1 promoter (Wils *et al*. 2010). Transgenic Thy1-hTDP-43 mice were generated by breeding male and female hemizygous TDP-43 mice. Genotyping for *TARDBP* was done by standard PCR on DNA phalanx biopsies, using primer sets designed by The Jackson Laboratory. Animals were bred and maintained at the Istituto di Ricerche Farmacologiche Mario Negri IRCCS, Milan, Italy, under standard conditions: temperature 21 ± 1 °C, relative humidity 55 ± 10%, 12h light schedule, and food and water *ad libitum*. For biochemical analysis, animals were deeply anesthetized with ketamine hydrochloride (IMALGENE, 150 mg/kg; cat. no. ALC104135013, Alcyon Italia) and medetomidine hydrochloride (DOMITOR, 2 mg/kg; cat. no. ALC100103011, Alcyon Italia) by intraperitoneal injection and perfused transcardially with 10 mL of phosphate-buffered saline (PBS). Mice were euthanized by exsanguination during transcardial perfusion under anesthesia. A total of 189 mice were used in this study. At the beginning of the experiment, the following numbers of animals were assigned to each genotype or treatment group: 21 non-transgenic and 59 Thy1-hTDP-43 mice for characterisation study; 48 Thy1-hTDP-43 mice treated with vehicle, 38 Thy1-hTDP-43 mice treated with SAHA, 6 Thy1-hTDP-43 mice treated with arimoclomol, 17 Thy1-hTDP-43 mice treated with SAHA+arimoclomol. The number of animals was calculated on the basis of experiments designed to reach a power of 0.8, with a minimum difference of 20% (α = 0.05).

### 2.7 Characterization of the Thy1-hTDP-43 mouse model by behavioral analysis

Thy1-hTDP-43 mice were characterized for disease progression by measuring body weight, hind limb extension reflex, gait impairment and tremor score. The extension reflex was quantified using the following 3-point scoring system: 3, hind limbs extending to an angle of 120 degrees; 2.5, hind limbs extending to <90 degrees with decreased reflex in one hind limb; 2.0, equal to 2.5 with decreased reflex in both hind limbs; 1. 5, loss of reflex with marked flexion in one hind limb; 1, equal to 1.5 with marked flexion in both hind limbs; 0.5, loss of reflex with hind limbs and paws held close to body but still able to walk; 0, equal to 0.5 but unable to walk (Pasetto *et al*. 2021). Gait impairment was scored from 0 to 3, as described (Zeballos C *et al*. 2023). In detail: 0 mice show no impairment; 1, mouse has a limp when walking; 2, mouse either has a severe limp or points its feet away from its body when walking; 3, mouse either has difficulty moving forward, shows minimal joint movement, does not use its feet to generate forward motion, has difficulty standing upright. Tremor was scored as described (Zeballos C *et al*. 2023): 0, no tremor observed; 1, mild tremor with movement; 2, severe tremor with movement; 3, severe tremor at rest and with movement. Onset of disease was determined retrospectively as the average age at which the mouse exhibited the first failure of maximum performance in the extension reflex score gait and tremor tests at two consecutive time points. Mice homozygous for TDP-43 in the C57BL/6J background exhibited overexpression of TDP-43 in neurons beginning at 7 days of age (pre-symptomatic stage), fast progressive motor behavioral impairment from 14±0.6 days of age (disease onset) and by 18±0.9 days of age developed severe symptoms (a score of 0.5 for the hind limb extension reflex or 3 for the gait or tremor tests). For behavioral analysis mice were euthanised through 30% CO_2_ inhalation when they reach a score of 0.5 for the hind limb extension reflex or 3 for the gait or tremor tests. We measured molecular biomarkers at 17 days of age (symptomatic stage), when the mice reach a score of 1 for the hind limb extension reflex and 2 for the gait or tremor tests (Supplementary Fig. 1).

### 2.8 Preclinical studies in the Thy1-hTDP-43 mouse model

Thy1-hTDP-43 mice received 50 mg/kg dose of SAHA, 10 mg/kg dose of arimoclomol (Arim.), 50 mg/kg dose of SAHA combined with 10 mg/kg dose of arimoclomol (SAHA+Arim.) or vehicle (Veh; 5% DMSO, 40% PEG300, 5% Tween-80, 50% Saline) through intraperitoneal injection, every three days, starting from 7 days of age. The impact of SAHA, or arimoclomol or SAHA combined with arimoclomol on protein biomarkers in Thy1-hTDP-43 mice was evaluated at disease onset (14 days of age for SAHA) or at symptomatic stage (17 days of age for SAHA, Arim. and SAHA+Arim.) through biochemical analysis. Furthermore, the impact of a 50 mg/kg dose of SAHA administered daily was assessed in Thy1-hTDP-43 mice from the age of 7 days until the age of 14 days. Mice were weighed daily, at the same time of day, on a balance with 0.1 g readability and ± 0.3 g linearity (cat. no. EU-C 7500PT, Gibertini). No exclusion criteria were pre-determined and no animals were excluded.

### 2.9 Subcellular fractionation in HEK293 cells, human PBMCs and mouse spinal cord

HEK293 cells were lysed in ice-cold RIPA buffer (0.3% Triton X-100, 50 mM Tris-HCl pH 7.4, 1 mM EDTA) and cOmplete™ Protease Inhibitor Cocktail Tablets (cat. no. 11836170001, Roche, 1 tablet/10 ml buffer), with rotation at 4 °C for 30 min, as previously described (Lauranzano *et al*. 2015). Cell extracts were centrifuged to pellet nuclei at 12000 x *g* for 10 min at 4 °C and the supernatant was collected in a new tube as cytoplasmic fraction. Nuclear pellet was washed by gently resuspension in ice-cold RIPA buffer and centrifuge at 10000 x *g* for 5 min at 4 °C. PBMCs were lysed essentially as described by Miskolci et al. (Miskolci *et al*. 2014). Cell pellets were suspended in ice-cold buffer containing 10 mM Hepes pH 7.5, 10 mM KCl, 3 mM NaCl, 3 mM MgCl_2_, 1 mM EDTA pH 8.8, 1 mM EGTA and cOmplete™ Protease Inhibitor Cocktail Tablets (cat. no. 11836170001, Roche, 1 tablet/10 ml buffer) and kept on ice for 15 min. Samples were treated by adding NP-40 at a final concentration of 0.5 %, vortexed and kept on ice for 8 sec, then centrifuged to pellet nuclei at 850 x *g* for 5 min at 4 °C, while the supernatant was collected in a new tube as cytoplasmic fraction. Nuclear pellet was resuspended in ice-cold RIPA buffer and centrifuge at 2400 x *g* for 5 min at 4 °C. Nuclear and cytoplasmic fractions were isolated from mouse spinal cord as described (Pasetto *et al*. 2021). Briefly, tissues were homogenized in six volumes (w/v) of buffer A (10 mM Tris-HCl pH 7.4, 5 mM MgCl_2_, 25 mM KCl, 0.25 M sucrose, 0.5 mM DTT) with cOmplete™ Protease Inhibitor Cocktail Tablets (cat. no. 11836170001, Roche, 1 tablet/10 ml buffer), and centrifuged at 800 x *g* for 10 min at 4 °C. The supernatant (cytoplasmic fraction) was centrifuged at 800 x *g* for 10 min at 4 °C to remove fraction contamination. The pellet was resuspended in buffer A, centrifuged at 800 x *g* for 10 min at 4 °C, subsequently resuspended in one volume of buffer A and one volume of buffer B (10 mM Tris-HCl pH 7.4, 5 mM MgCl_2_, 25 mM KCl, 2 M sucrose, cOmplete™ Protease Inhibitor Cocktail Tablets (cat. no. 11836170001, Roche, 1 tablet/10 ml buffer) and loaded on a layer of one volume of buffer B. Samples were ultracentrifuged at 100000 x *g* for 45 min at 4 °C and the pellet (nuclear fraction) was resuspended in 100 μL of buffer A, centrifuged at 800 x *g* for 10 min at 4 °C and resuspended in 40 μL buffer A. For HEK293 cells, human PBMCs and mouse spinal cord, cytoplasmic and nuclear fractions were lysed by adding pre-warmed at 95 °C SDS at a final concentration of 1% and shaken at 800 rpm for 5 min at 95 °C. Nuclei were then sonicated for 30 sec at RT. GAPDH and laminin A/C were used as cytoplasmic and nuclear markers, respectively, for the experiments with HEK293 cells, PBMCs and mouse spinal cord, as shown in Supplementary Fig. 2. Given the abundance of nuclear TDP-43, the analysis of nuclear and cytoplasmic fractions was done separately throughout the study to avoid signal saturation and to allow detection of cytoplasmic changes.

### 2.10 Total protein extraction in HEK293 cells, human PBMCs, mouse spinal cord and sciatic nerve

HEK293 cells, human PBMCs or lumbar spinal cord were lysed in pre-warmed at 95 °C buffer containing 1% SDS, cOmplete™ Protease Inhibitor Cocktail Tablets (cat. no. 11836170001, Roche), shaken at 800 rpm for 5 min at 95 °C in the Thermomixer and DNA complexes were disrupted using a 25-gauge needle of a 1 ml syringe. Samples were centrifuged at 9000 x *g* for 15 min at RT and supernatant was collected in a new tube as total protein extract. The extraction of total protein from sciatic nerves was carried out as reported (Tortarolo *et al*. 2015). Briefly, sciatic nerve was homogenized in ice-cold lysis buffer (50 mM Tris-HCl pH 8, 150 mM NaCl, 5 mM EGTA, 2 mM MgCl_2_, 1% of Triton X-100, 10% anhydrous glycerol, protease and phosphatase inhibitors), sonicated, centrifuged at 18000 x *g* for 15 min at 4 °C and supernatant was collected as total protein extract.

### 2.11 Extraction of detergent-insoluble proteins

Spinal cord was homogenized in 10 volumes (w/v) of buffer (15 mM Tris–HCl pH 7.6, 1 mM dithiothreitol, 0.25 M sucrose, 1 mM MgCl_2_, 2.5 mM EDTA, 1 mM EGTA, 0.25 M Na_3_VO_4_, 2 mM Na_4_P_2_O_7_, 25 mM NaF, 5 µM MG132, cOmplete™ Protease Inhibitor Cocktail Tablets (cat. no. 11836170001, Roche, 1 tablet/10 ml buffer) as described (Pasetto *et al*. 2021). Briefly, the samples were centrifuged at 10000 x *g*, and the pellet was suspended in an ice-cold homogenization buffer containing 2% Triton X-100 and 150 mM KCl. The samples were then centrifuged at 10000 x *g* to obtain the Triton-insoluble (insoluble) fraction, finally suspended in buffer containing 2% SDS.

### 2.12 Dot blot and Western Blot (WB)

Protein levels were determined using the Pierce™ BCA Protein Assay Kit (cat. no. 23227, Thermo Fisher Scientific). For WB, samples (15-20 µg) were separated in 12% SDS-polyacrylamide gels and transferred to polyvinylidene difluoride membranes (0.45 µm; cat. no. 1620115, Millipore), as described previously (Basso *et al*. 2009). Dot blot was used after verification that the antibody detected only specific bands in WB, as described (Filareti *et al*. 2017). Proteins (3 µg) were loaded directly onto nitrocellulose Trans-blot transfer membranes (0.45 µm; cat. no. 1620115, Bio-Rad), depositing each sample on the membrane by vacuum filtration, as described previously (Luotti *et al*. 2020). WB and dot blot membranes were blocked with 3% (w/v) BSA (cat. no. 7906-100G, Sigma-Aldrich) and 0.1% (v/v) Tween 20 in Tris-buffered saline, pH 7.5, and incubated with primary antibodies and then with peroxidase-conjugated secondary antibodies (cat. no. 171193520 anti-mouse, cat. no. 17376631 anti-rabbit, GE Healthcare). Blots were developed with the Luminata Forte Western Chemiluminescent HRP Substrate (cat. no. WBLUF0500, Millipore) on the ChemiDoc™ Imaging System (Bio-Rad). Densitometry was done with Image Lab 6.0 software (Bio-Rad). The immune reactivity of the different proteins was normalized to Ponceau Red staining (cat. no. 09189-1L-F, Fluka) and, in some cases, also to an internal standard (IS) to compare samples from the same experiment loaded in different immunoblot. Uncropped images of the WB are included in the Supplementary material.

### 2.13 Capillary electrophoresis immunoassay (CEI)

CEI was performed on the Jess instrument (Protein Simple, Biotechne). Samples diluted in 0.1X Sample Buffer to 0.2-2 µg/µl concentrations, added to the Fluorescent Master Mix and incubated for 5 min at 95 °C. Samples, antibodies and reagents were loaded in the plate following the “Immunoassay + Total Protein” template and plate was centrifuged at 1000 x *g* for 5 min at RT, according to the manufacturer’s instructions. 12–230 kDa separation module with 25-capillary cartridge was employed and chemiluminescent detection with RePlex and Total Protein assay was run. Target proteins are normally observed to be approximately 10 kDa higher than in standard WB, as described (Fourier *et al*. 2019). Data were analysed using Compass for Simple Western 6.3.0 software and immunoassay data were corrected for total protein signal.

### 2.14 Generation and purification of an anti-K125 acetylated PPIA (PPIAK125ac) polyclonal antibody

For the preparation of polyclonal antibodies, the peptide WLDGKHVVFG acetylated at the K residue was synthesized by solid-phase peptide synthesis on an automated Alstra synthesizer (Biotage, Uppsala, Sweden) at 0.1 mM scale on tetravalent multiple antigenic peptide resin (MAP) resins using Fmoc-protected L-amino acids (Sigma Aldrich, Laufelfingen, CH) (Kowalczyk *et al*. 2011). Amino acids were activated by a reaction with O-(Benzotriazole-1-yl)-N, N, N’, N’-tetramethyluronium tetrafluoroborate, and N, N-diisopropylethylamine. A capping step with acetic anhydride was included after the last coupling cycle of each amino acid. The peptide was cleaved from the resin with trifluoroacetic acid/thioanisole/water/ phenol/ethanedithiol (82.5:5:5:5:2.5 vol/vol), precipitated, and washed with diethyl ether. The peptide was purified by reverse-phase high-performance liquid chromatography on a semi-preparative C18 column (Waters Corporation, Milford, MA), and its identity was confirmed with a MALDI-TOF spectrometer (Applied Biosystems, Concord, Ontario, Canada). The peptide purity was higher than 95%. The samples were then freeze-dried and stored at −20 °C until use. The polyclonal antibodies production was carried out by the Company Biogem, Ariano Irpino, Avellino (Italy). Immunization was performed by subcutaneous injection of the MAP peptide in White New Zealand rabbits, and the titer was measured by ELISA on serum samples. A highly specific antibody for the acetylated form of PPIA was obtained through a two-step affinity purification process using the Pierce™ NHS-Activated Agarose kit (cat. no. 26198). Initially, the serum was deprived of antibodies that recognize the peptide corresponding to the amino acids of PPIA from position 121 to 130, but whose K125 was not acetylated. The resulting serum was then further affinity-purified, performing positive selection by resin functionalization with the same peptide used for rabbit immunization, acetylated at K125. To verify specificity, the purified antibody was tested by dot blot against a range of peptide amounts (0.03 to 4 μg), both acetylated and non-acetylated (Supplementary Fig. 3).

### 2.15 Mutant PPIA transfection

HEK293 cells were transfected with plasmids containing PPIA WT, PPIA K125Q or PPIA K125R Myc-tagged constructs, previously generated as described (Lauranzano *et al*. 2015). In detail, 70% confluent HEK293 cells were transfected with 0.016 μg/μL plasmid solution complexed with Lipofectamine™ 2000 Reagent (cat. no. 11668-027, Invitrogen) in Opti-MEM™ I Reduced Serum Medium (cat. no. 31985-070, Gibco), then incubated at 37 °C with humidified atmosphere in a CO_2_ incubator for 48 hours and lysed for detergent-insoluble protein extraction as previously described (Luotti *et al*. 2020).

### 2.16 HDAC mRNA silencing

For RNA interference experiments in HEK293 cells, a reverse transfection protocol was applied. Complexes were prepared in 24-wells plate by diluting 6 pmol of siRNA for *HDAC1* (5’-UGG CCA UCC UGG AAC UGC UAA AGU A-3’; ID: HSS104726; cat. no. 1299001, Invitrogen) or siRNA for *HDAC3* (5’-CAA GGA AAG CGA UGU GGA GAU UUA A-3’; ID: HSS189614; cat. no. 1299001, Invitrogen) or scrambled siRNA (Stealth™ RNAi Negative Control Duplex cat. no. 12935-113; Invitrogen) in Opti-MEM™ I Reduced Serum Medium (cat. no. 31985-070, Gibco) with Lipofectamine™ RNAiMAX Transfection Reagent (cat. no. 13778030, Invitrogen). After a 15 min incubation at RT, cells were diluted in complete growth medium without antibiotics, added to each well and incubated for 24 hours at 37 °C in a humidified atmosphere and 5% CO_2_, according to the manufacturer’s protocol. Gene silencing was confirmed by immunoblot analysis.

### 2.17 Immunohistochemistry

For immunohistochemical analysis, animals were deeply anesthetized with ketamine hydrochloride (IMALGENE, 150 mg/kg; cat. no. ALC104135013, Alcyon Italia) and medetomidine hydrochloride (DOMITOR, 2 mg/kg; cat. no. ALC100103011, Alcyon Italia) by intraperitoneal injection, perfused transcardially with 10 mL of phosphate-buffered saline (PBS) followed by 100 mL of 4% paraformaldehyde (cat. no. 1.04002.1000, Merck) in PBS. Mice were euthanized by exsanguination during transcardial perfusion under anesthesia. Spinal cord was rapidly removed, postfixed for 3h, transferred to 20% sucrose in PBS overnight and then to 30% sucrose solution until they sank, frozen in N-pentane at −45 °C and stored at −80 °C. Before freezing, spinal cord was divided into cervical, thoracic, and lumbar segments and included in Tissue-tec OCT compound (cat. no. 4583, Sakura). Coronal sections (30 µm; four slices per mouse) of lumbar spinal cord were then sliced and immunohistochemistry was done. Rat monoclonal anti-CD11b antibody (1:800, Bio-Rad; RRID: AB_ 321292) was used for immunohistochemistry analysis. Briefly, slices were incubated for 1h at RT with blocking solutions (0.3% Triton X100 plus 10% NGS), then overnight at 4 °C with the primary antibodies. After incubation with biotinylated secondary antibody (1:200; 1h at RT; Vector Laboratories) immunostaining was developed using the avidin–biotin kit (Vector Laboratories) and diaminobenzidine (cat. no. K3468, Sigma). Coronal lumbar spinal cord (30 µm; twelve slices per mouse) were stained with 0.5% cresyl violet to detect the Nissl substance of neuronal cells. Stained sections were collected at 20 X and 40 X with an Olympus BX-61 Virtual Stage microscope so as to have complete stitching of the whole section, with a pixel size of 0.346 µm. Acquisition was done over 6-µm-thick stacks with a step size of 2 µm. The different focal planes were merged into a single stack by mean intensity projection to ensure consistent focus throughout the sample. Finally, signals were analysed for each slice with ImageJ and OlyVIA software.

### 2.18 Plasma neurofilament light chain (NfL) and matrix metalloproteinase-9 (MMP-9) quantification

Mouse plasma samples were collected in K2-EDTA (cat. no. 367862, BD) Microtainer blood collection tubes and centrifuged at 5000 x *g* for 5 min to obtain the plasma. The NfL was quantified using a Simoa NfLight Advantage (SR-X) Kit (cat. no. 103400) on a Quanterix SR-X platform (Quanterix, Boston, MA, USA). The level of MMP-9 in plasma was measured with an AlphaLISA kit for the murine protein (cat. no. AL519, Revvity). AlphaLISA signals were measured using an Ensight Multimode Plate Reader (PerkinElmer). All reagents used for NfL and MMP-9 analysis were from a single lot, and measurements were performed according to the manufacturer’s protocol.

### 2.19 Quantitative real-time polymerase chain reaction

Total RNA from gastrocnemius muscle and spinal cord was extracted using Trizol (cat. no. 15596026, Invitrogen, Waltham, MA, USA) and purified with PureLink RNA columns (cat. no. 12183018A, Life Technologies, Carlsbad, CA, USA). RNA samples were treated with DNase I and reverse transcribed with the High-Capacity cDNA Reverse Transcription Kit (cat. no. 4368814, Life Technologies, Carlsbad, CA, USA). For quantitative real-time polymerase chain reactions (RT-PCR), we used the Taq Man Gene expression assay (Applied Biosystems, Waltham, MA. USA), on cDNA specimens in triplicate, using SensiFAST Probe Lo-ROX master mix (cat. no. BIO-84005, Meridian Bioscience) and 1X mix containing specific receptor probes for mouse acetylcholine receptor (*AChR*) γ-subunit in muscle (Mm00437419_m1; Life Technologies, Carlsbad, CA, USA), microtubule associated protein 2 (*Map-2*) (Mm00485231_m1; Life Technologies, Carlsbad, CA, USA), *Tardbp* (cat. no. Mm01257504_g1; Life Technologies, Carlsbad, CA, USA), *Ppia* (cat. no. Mm02342430_g1; Life Technologies, Carlsbad, CA, USA), *Hdac1* (cat. no *Mm02745760*_g1; Life Technologies, Carlsbad, CA, USA), *Hdac2* (cat. no Mm00515108_m1; Life Technologies, Carlsbad, CA, USA), *Hdac3* (cat. no. Mm00515916_m1; Life Technologies, Carlsbad, CA, USA), *Hdac4* (cat. no. Mm01299557_m1; Life Technologies, Carlsbad, CA, USA) and *Hdac6* (cat. no. Mm00515945_m1; Life Technologies, Carlsbad, CA, USA) in spinal cord. Relative quantification was calculated from the ratio of the cycle number (Ct) at which the signal crossed a threshold set within the logarithmic phase of the given gene to that of the reference mouse β*-actin* gene (cat. no Mm02619580_g1; Life Technologies, Carlsbad, CA, USA). The means of the triplicate results for each sample were used as individual data for 2^−ΔΔCt^ statistical analysis.

### 2.20 Skiptic and cryptic exon analysis

Total RNA from spinal cord was extracted using Trizol (cat. no. 15596026, Invitrogen, Waltham, MA, USA) and purified with PureLink RNA columns (cat. no. 12183018A, Life Technologies, Carlsbad, CA, USA). RNA samples were treated with DNase I and reverse transcribed with the High-Capacity cDNA Reverse Transcription Kit (cat. no. 4368814, Life Technologies, Carlsbad, CA, USA). For the evaluation of skiptic exon Ankrd42, Ube3C, Slc6a6, Plod1, Pex16, Pacrgl, Herc2 and of cryptic exon 2610507B11Rik, Adipor2, A230046K03Rik, Adnp2, Celf5 in spinal cord semi-quantitative PCR was performed on cDNA with specific primers spanning the differentially expressed exon. Products were electrophoresed on 8% polyacrylamide gel and using GelRed (cat. no. SCT123, Merck Millipore) staining results were visualized on a ChemiDoc Imaging System (Bio-Rad, Hercules, CA, USA). Results were analysed using the percentage of exon exclusion (PEE) measure, calculated dividing the intensity of the exon exclusion band by the sum of both exon inclusion and exon exclusion bands. Primer sequences are reported in Supplementary Table 1.

### 2.21 Blinding and randomization procedures

For the animal studies (behavior and treatments), a blinding approach was employed, entailing the utilisation of neutral identifiers for mice and the delineation of distinct roles among researchers. The experimenter was unaware of study groups during treatment and subsequent biochemical analysis. The randomization process was conducted using a block randomization approach, whereby mice were assigned randomly to different groups of treatment. The responsibility for assigning the mice to the various experimental groups and providing the compounds for treatment, with the compounds being defined as Compound A and B, without defining which is vehicle and which is drug compound, fell to one of the researchers (R1). The two other researchers (R2-R3) treated the animals as instructed by R1 and conducted the behavioral tests, collected the tissues and performed the biomarker analysis. When all experiments and outcomes were collected, R1 opened the blind to reveal the treated and vehicle groups. For all other experiments in cell models, blinding procedures were also applied, as data analysis was conducted by a researcher different from the experimenter.

### 2.22 Statistical analysis

Prism 10.0 (GraphPad Software Inc., San Diego, CA) was used. Normality was assessed using the Shapiro–Wilk test. For each variable, the differences between experimental groups were analysed by one- or two-tailed unpaired t test, one- or two-tailed paired t test, one-way ANOVA followed by post-hoc tests when normally distributed or by one- or two-tailed Mann-Whitney test, Kruskal-Wallis test when no normally distributed. Two-way ANOVA for repeated measure followed by Sidak’s or Tukey’s post-hoc test or multiple Mann-Whitney tests was used to analyse body weight loss and behavioral analysis in Thy1-hTDP-43 mice respectively when normally or not distributed. Log-rank (Mantel-Cox) test was used for onset and survival evaluation in Thy1-hTDP-43 mice. P values below or equal to 0.05 were considered significant. No test for outliers was conducted. Sample sizes were determined a priori based on prior published studies (Lauranzano *et al*. 2015; Pasetto *et al*. 2021; Caldi Gomes *et al*. 2024) to reach a power of 0.8, with a minimum difference of 20% (α = 0.05).

## 3 Results

### 3.1 SAHA reverses TDP-43 mislocalization and increases PPIA acetylation in an *in vitro* model

To investigate the role of Lys-acetylation on TDP-43 proteinopathy we tested a number of HDACi on a cellular model of chronic nutrient starvation (Reineke *et al*. 2018). Under these conditions, HEK293 cells present cytoplasmic mislocalization of TDP-43 and its C-terminal fragment of 35 kDa (TDP-35CTF) (Fig. 1A-B and Supplemenatary Fig. 4A). We found that in our assay SAHA was the most effective in restoring control conditions by analyzing both full-length TDP-43 and TDP-35CTF, and this effect was dose-dependent (Fig. 1A-B and Supplementary Fig. 4A-C).

**FIGURE 1.**
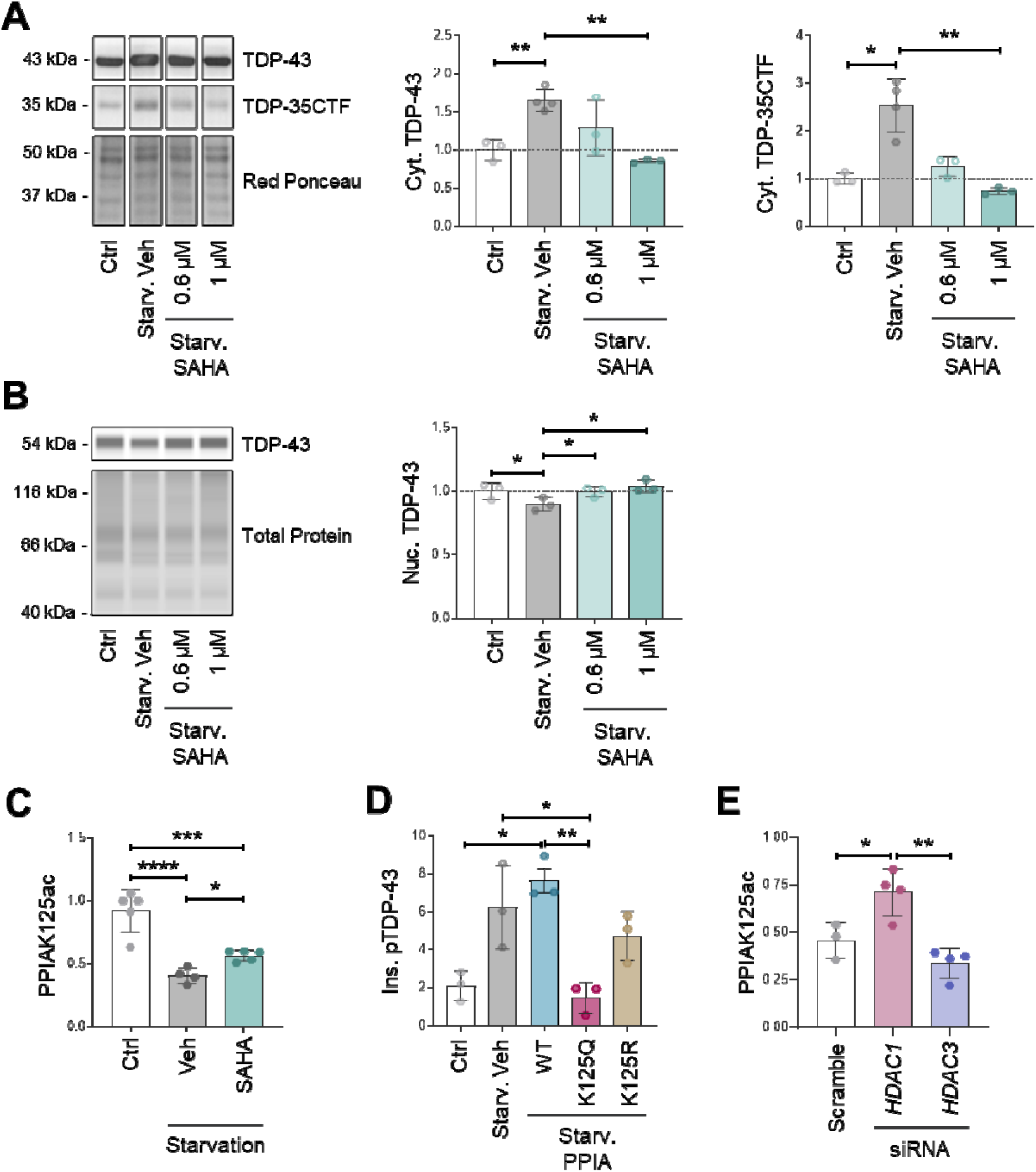
SAHA reduces TDP-43 proteinopathy and increases acetyl-PPIA in an *in vitro* model. (**A**) Cytoplasmic (Cyt.) levels of TDP-43 and TDP-35CTF in control (Ctrl; n=3) and serum-deprived HEK293 cells (Starv.), overnight pre-treated with vehicle (Veh; n=4) or with 0.6 and 1 μM SAHA (n=3). Representative WB is shown and corresponding uncropped blot is reported in Supplementary Fig. 4A. (**B**) Nuclear (Nuc.) levels of TDP-43 in control (Ctrl; n=3) and serum-deprived HEK293 cells (Starv.), overnight pre-treated with vehicle (Veh; n=3) or with 0.6 and 1 μM SAHA (n=3). CEI’s representative virtual lane view is reported. (**A-B**) Dashed line indicates Ctrl level and treated conditions were normalized to Ctrl level. (**C**) PPIAK125ac levels in total lysate of control (Ctrl; n=5) and serum-deprived HEK293 cells (Starvation), overnight pre-treated with vehicle (Veh; n=5) or with 1 μM SAHA (n=6). (**D**) Insoluble (Ins.) pTDP-43 levels in control (Ctrl; n=3) and serum-deprived HEK293 cells (Starv.), vehicle-treated (Veh; n=3) or transiently transfected for 48 hours with PPIA K125Q (n=3), PPIA K125R (n=3) mutants, or PPIA WT (n=3). (**E**) PPIAK125ac levels in total lysate of HEK293 transiently transfected for 24 hours with siRNA control (Scramble; n=3), siRNA for HDAC1 (n=4) and HDAC3 (n=4). (**A-E**) In all experiments n refers to the number of independent cell culture preparations. Data (mean ± SD) indicate WB (**A,C,E**), CEI (**B**) or dot blot (**D**) immunoreactivity normalized to total protein loading (**A-E**) and to PPIA levels (**C,E**). *p ≤ 0.05, **p < 0.01, ***p < 0.001, ****p < 0.0001, by one-way ANOVA, Dunnett’s multiple comparisons (**A,** Cyt. TDP-43), Uncorrected Fisher’s LSD (**B-C**) or Tukey’s multiple comparisons (**E**) post-hoc tests, or by Kruskal-Wallis test, Uncorrected Dunn’s multiple comparisons test (**A**, Cyt. TDP-35CTF,**D**).

To evaluate the effect of SAHA on PPIA acetylation (PPIAK125ac) we developed an antibody against a stretch of the PPIA sequence centered on the acetylated K125 (Supplementary Fig. 3). We found that under chronic nutrient starvation conditions there was a substantial decrease in PPIAK125ac and that SAHA was able to significantly increase it (Fig. 1C). On the other hand, serum deprived HEK293 cells transfected with the acetylation-mimetic mutant PPIA K125Q displayed low levels of pTDP-43 in the insoluble fraction compared with cells transfected with PPIA WT or the acetylated-defective mimetic PPIA K125R mutant (Fig. 1D), which poorly interacts with TDP-43 (Lauranzano *et al*. 2015). Therefore, we can hypothesize that deacetylation of PPIA under stress conditions may impair PPIA chaperone activity toward TDP-43. We confirmed by WB that all PPIA constructs were expressed at similar levels in the cells (Supplementary Fig. 4F).

SAHA has a relatively broad specificity, but is a potent inhibitor of HDAC1 and HDAC3 that are highly expressed in the nucleus (El-Awady *et al*. 2021). By knocking down experiments, we could deduce that HDAC1, and not HDAC3, deacetylates PPIAK125ac (Fig. 1E and Supplementary Fig. 4D-E), however we cannot exclude that other HDACs could be involved.

We concluded that SAHA is an interesting candidate to mitigate TDP-43 proteinopathy potentially through the modulation of PPIAK125 acetylation. We therefore extended our investigation to assess the effect of SAHA on TDP-43and PPIA acetylation in PBMCs of ALS patients and in a mouse model of TDP-43 proteinopathy.

### 3.2 SAHA reduces cytoplasmic TDP-43 by increasing PPIAK125ac in PBMCs of ALS patients

Previous studies detected TDP-43 mislocalization in PBMCs of ALS and ALS-FTD patients (De Marco *et al*. 2011). We tested whether treatment with SAHA can affect TDP-43 localization in PBMCs of ALS patients. We selected 9 ALS patients with PBMCs displaying accumulation of TDP-43 and TDP-35CTF in the cytoplasm, as detected by CEI, and low PPIAK125ac in the nucleus compared to age- and sex-matched healthy controls (Fig. 2A-E and Table 1).

**FIGURE 2.**
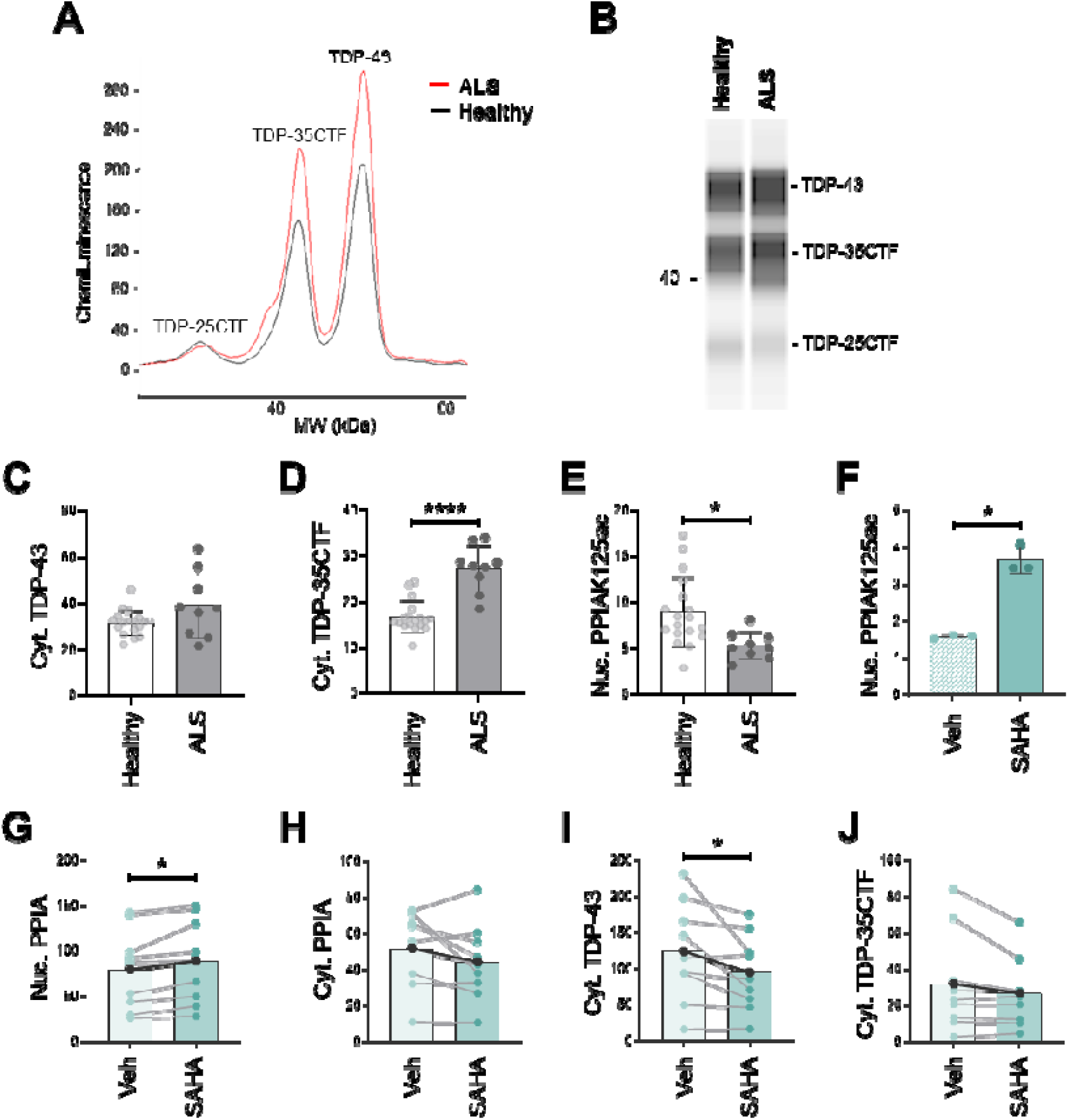
SAHA increases PPIAK125ac and reduces cytoplasmic TDP-43 in PBMCs of ALS patients. Electropherogram (**A**) and virtual lane view (**B**) of the TDP-43 signal by CEI in cytoplasmic (Cyt.) fractions of PBMCs from an ALS patient (red line) and a healthy subject (black line). Peaks for TDP-43, TDP-35CTF and TDP-25CTF are indicated. (**C-E**) CEI analysis of cytoplasmic TDP-43 and TDP-35CTF (**C**-**D**), and nuclear (Nuc.) PPIAK125ac (**E**) in PBMCs of ALS patients (n=9) and healthy subjects (n=18). Data (mean ± SD) indicate CEI immunoreactivity normalized to total protein (**C-E**) and to PPIA levels (**E**). *p < 0.05, ****p < 0.0001 by Mann Whitney test (**D**) or unpaired t test (**E**). (**F-J**) CEI analysis of nuclear PPIAK125ac (**F**), nuclear and cytoplasmic PPIA **(G-H)**, and cytoplasmic TDP-43 and TDP-35CTF (**I**-**J**) in the cultured PBMCs of the same ALS patients overnight treated with vehicle (Veh) or with 1 μM SAHA. (**F**) Data (mean ± SD; n=3 technical replicates derived from pools of the 9 ALS patients Veh or SAHA treated) indicate CEI immunoreactivity normalized to total protein and to PPIA levels. *p ≤ 0.05 by Mann-Whitney test. (**G-J**) Data (mean ± SD; n=9 independent PBMC culture preparations from ALS patients) are shown as slope chart and indicate CEI immunoreactivity of overnight SAHA-treated PBMCs, normalized to total protein loading, in comparison with the relative vehicle-treated sample. The black slope chart indicates the mean value for each protein target. *p < 0.05 by paired t test (**G,I**).

Two patients carried a TDP-43 mutation (TDP-43^G295S^, TDP-43^S393L^), but their TDP-43 distribution in PBMCs did not differ from the other sporadic patients. PBMCs from each patient were treated or not with 1 µM SAHA overnight. SAHA increased PPIAK125ac, concomitantly induced nucleus-cytoplasmic redistribution of PPIA (Fig. 2F-H), along with a reduction in cytoplasmic TDP-43 and TDP-35CTF levels, although the latter did not reach statistical significance (Fig. 2I–J). This occurred without a noticeable increase of TDP-43 in the nucleus, where it is already highly concentrated (data not shown). The greater effect on full-length TDP-43 is likely due to the presence of a nuclear localization signal, which is absent in TDP-35CTF. These findings suggest that acetylated PPIA may contribute to the regulation of TDP-43 localization and support the possible use of PBMCs as a platform for evaluating drugs targeting TDP-43 mislocalization.

### 3.3 Homozygous Thy1-hTDP-43 mouse is a severe model of TDP-43 proteinopathy

To test the effect of SAHA *in vivo*, we used the homozygous Thy1-hTDP-43 mouse model of ALS, which develops marked TDP-43 pathology and severe quadriplegia (Wils *et al*. 2010). The Thy1 promoter drives TDP-43 expression starting at postnatal day 7, enabling the study of the mechanisms leading to TDP-43 proteinopathy *in vivo* from a defined onset. To reduce phenotypic variability, the original B6;SJL hybrid strain was backcrossed to a fully congenic C57BL/6J strain. This resulted in a highly reproducible onset of symptoms at 14 days of age (Supplementary Fig. 1). However, unexpectedly, the disease became even more aggressive, limiting behavioral analysis and studies on disease progression.

To evaluate the effect of drugs in proof-of principle studies, we identified a number of molecular biomarkers as outcome measures up to a symptomatic stage (17 days). Thy1-hTDP-43 mice show a dramatic loss of motor neurons in the spinal cord already at the onset of the disease mirrored by a great increase in plasmatic NfL (Fig. 3A-B). Neurodegeneration is accompanied by microgliosis that becomes significant respect to control at the symptomatic stage (Fig. 3C). In the spinal cord, the TDP-43 in the nucleus greatly decreases as disease progresses and accumulates in the cytoplasm, especially as TDP-35CTF and as detergent-insoluble phosphorylated form (Fig. 3D-F). To evaluate the effects of TDP-43 pathology on splicing in the spinal cord of the Thy1-hTDP-43 mouse, we evaluated the exclusion or the inclusion respectively of a number of skiptic and cryptic exons, previously identified in mutant TDP-43 mouse models (Fratta *et al*. 2018). All of the skiptic exons evaluated (Herc2, Ankrd42, Ube3C, Slc6a6, Plod1, Pex16, Pacrgl) were detected and the level of the exon exclusion increased as disease progresses, particularly for Ankrd42, Slc6a6 and Herc2 that already at the onset were significantly higher respect to the presymptomatic stage (Fig. 3G-J and Supplementary Fig. 5A-D). Instead, none of the cryptic exons analyzed (Adipor2, Adnp2, A230046K03Rik, 2610507B11Rik, Celf5) were detected (data not shown). PPIA essentially followed TDP-43, translocating to the cytoplasm as disease progresses possibly recruited in membrane-less assemblies and/or aggregates (Fig. 3K), while PPIAK125ac is retrieved to the nucleus, probably recruited into nuclear bodies as a response to stress (Fig. 3L) (Wang *et al*. 2020). Consequently, there is a loss of PPIA in the sciatic nerve (Fig. 3M), where also high levels of pTDP-43 are detected (Fig. 3N). The sciatic nerve shows a great decrease in acetylated tubulin at the onset of the disease (Fig. 3O), suggesting an early impairment of the axonal transport. In the gastrocnemius muscle of the Thy1-hTDP-43 mice, the expression of fetal *AChR* γ-subunit, which in mice is normally downregulated within the first two weeks after birth (Missias *et al*. 1996), substantially increases at a symptomatic stage compared to the onset (Fig. 3P), highlighting muscle denervation, as previously observed (Margotta *et al*. 2023).

**FIGURE 3.**
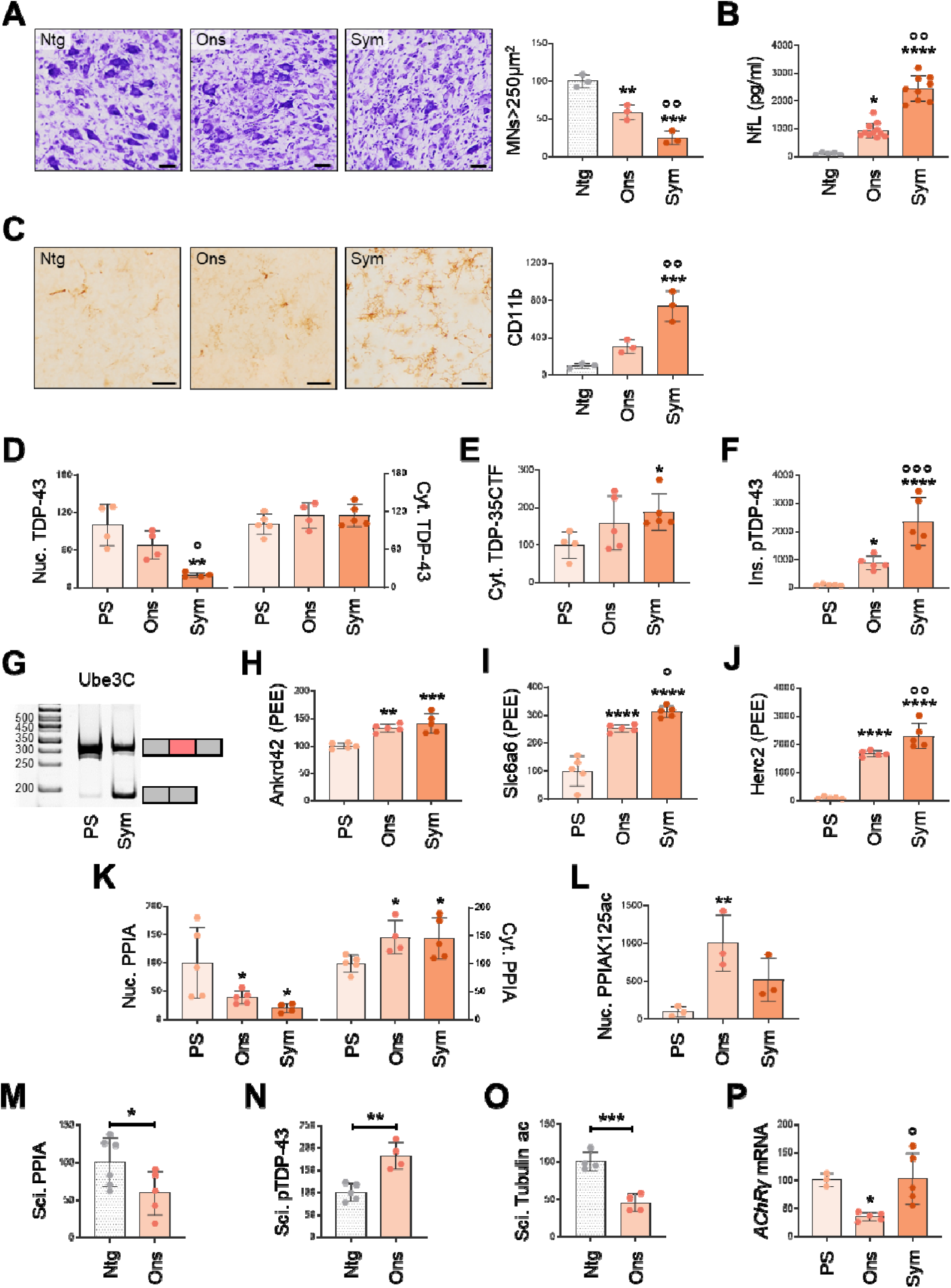
Homozygous Thy1-hTDP-43 mouse is a severe model of TDP-43 proteinopathy. (**A-Q**) Thy-hTDP-43 mice were characterized at pre-symptomatic (PS), disease onset (Ons) and symptomatic (Sym) stages for ALS pathological markers. (**A,C**) Quantification of Nissl-stained motor neurons (MNs > 250 μm^2^) and CD11b-stained microglia in lumbar spinal cord hemisections from non-transgenic (Ntg) and Thy1-hTDP-43 mice at Ons and Sym stage. Data are mean ± SD (n=3 mice/group). **p < 0.01 and ***p < 0.001 versus Ntg, °°p < 0.01 versus Ons by one-way ANOVA, Tukey’s multiple comparisons. Representative Nissl- and CD11b-stained lumbar spinal cord sections are shown. Scale bar 50 µm. (**B**) NfL plasma levels in Ntg mice (n=5) and Thy1-hTDP-43 mice at Ons (n=11) and Sym stages (n=9). Data are mean ± SD. *p < 0.05 and ****p < 0.0001 versus Ntg, °°p < 0.001 versus Ons by Kruskal-Wallis test, Uncorrected Dunn’s comparisons test. (**D-F**) Analysis of nuclear (Nuc.) and cytoplasmic (Cyt.) TDP-43 (**D**), cytoplasmic TDP-35CTF (**E**) (WB imagine in Supplementary Fig. 9A-B) and insoluble (Ins.) pTDP-43 (**F)** in spinal cord of Thy1-hTDP-43 mice at PS, Ons and Sym stages. Data (mean ± SD; n=4-5 mice/group) indicates immunoreactivity by dot blot (**D,F**) or WB (**E**) normalized to total protein loading. *p < 0.05, **p < 0.01 and ****p < 0.0001 versus PS, °p < 0.05 and °°°p < 0.001 versus Ons by one-way ANOVA, Tukey’s multiple comparisons (**D**) or Uncorrected Fisher’s LSD (**F**) post hoc test, by Mann Whitney test (**E**). (**G**) Diagram illustrating skiptic exon (right) and representative lanes of Ube3C skiptic exon from acrylamide gel in spinal cord of Thy1-hTDP-43 mice at PS and Sym stages (left). **(H-J)** Semi quantitative PCR of Ankrd42 (**H**), Slc6a6 (**I**) and Herc2 (**J**) skiptic exons in spinal cord of Thy1-hTDP-43 mice at PS, Ons and Sym stages. Data (mean ± SD; n =5 mice/group) are expressed as PEE. **p < 0.01, ***p < 0.001 and ****p < 0.0001 versus PS, °p < 0.05 and °°p < 0.01 versus Ons by one-way ANOVA, Tukey’s multiple comparisons (**H-J**). (**K-L**) Analysis of nuclear and cytoplasmic PPIA (**K**) and nuclear PPIAK125ac (**L**) in spinal cord of Thy1-hTDP-43 mice at PS, Ons and Sym stages. Data (mean ± SD; n=3-5 mice/group) indicate immunoreactivity by dot blot (**K**) or CEI (**L**) normalized to total protein loading (**K-L**) and to PPIA (**L**) levels. *p < 0.05, **p < 0.01 versus PS by one-way ANOVA, Uncorrected Fisher’s LSD post-hoc test. (**M-O**) Analysis of PPIA (**M**), pTDP-43 (**N**) and acetyl tubulin (Tubulin ac) **(O)** in sciatic nerve (Sci.) of Ntg mice (n=4-6) and Thy1-hTDP-43 mice at Ons (n=4-5). Data (mean ± SD) indicate dot blot immunoreactivity normalized to total protein loading (**M-O**) and to tubulin (**O**) levels. *p < 0.05, **p < 0.01, ***p < 0.001 by unpaired t test. (**P**) RT-PCR for AChR γ-subunit in gastrocnemius muscle of Thy1-hTDP-43 mice at PS, Ons and Sym stages (n=3-5 mice/group). Data (mean ± SD) are normalized to β-actin and reported as relative mRNA expression. *p < 0.05 versus PS, ° p < 0.05 versus Ons by one-way ANOVA with Tukey’s multiple comparisons. (**A,C-P**) Data are expressed as percentage of Ntg mice (**A,C,M-O**) or Thy1-hTDP-43 mice at PS stage of disease (**D-F, H-L,P**).

### 3.4 SAHA delayed the onset of TDP-43 pathology and in combination with arimoclomol reduced neuromuscular pathology markers in the mouse model

We established optimal treatment schedule for SAHA by a pilot study in which Thy1-hTDP-43 mice were treated intraperitoneally with 50 mg/kg every day or every three days from presymptomatic stage up to 14 days of age (Supplementary Fig. 6). The less frequent treatment schedule was more effective, on the basis of lower plasmatic NfL concentration and TDP-35CTF accumulation in the cytoplasm (Supplementary Fig. 6C-D) and therefore was chosen for the following trials. We performed two studies with two independent groups of mice with the aim to establish if SAHA interfered with the first phases of the pathology and if the effect was maintained over time.

In the first study, mice were treated up to 14 days of age (onset of the disease) (n=17 vehicle, n=18 SAHA) and sacrificed to evaluate the effect of the treatment at a molecular level using the previously established outcome measures (Fig. 4A). Compared to vehicle-treated, SAHA-treated mice had lower NfL concentration in plasma, higher *Map*-2 expression in spinal cord and lower MMP-9 in plasma (Fig. 4B-D), indexes of a reduced neurodegeneration and neuroinflammation. TDP-43 was higher in the nucleus and TDP-35CTF accumulation was lower in the cytoplasm (Fig. 4E-F) and there was a tendency to a lower pTDP-43 aggregation (Fig. 4G). A milder TDP-43 pathology was associated with a significant decrease in the exclusion of the skiptic exons for Plod1, Pex16, Herc2 and Ube3C (Fig. 4H-K), a reduction in the nuclear translocation of PPIAK125ac (Fig. 4L), that is high at the onset of the disease (Fig. 3L) and likely a marker of stress response, no reduction of nuclear PPIA and a higher level of PPIA in the cytoplasm (Fig. 4M). In periphery, possibly as a consequence of a reduced PPIA recruitment in membrane-less assemblies or aggregates, we detected an increased level of PPIA in the sciatic nerve (Fig. 4N) and decreased pTDP-43 (Fig. 4O). We also detected increased acetylated tubulin (Fig. 4P) and decreased *AChR* γ-subunit expression in the gastrocnemius (Fig. 4Q), suggesting a delayed impairment in the neuromuscular function. Finally, we also checked the effect of SAHA on *Hdac1, 2, 3, 4*, and *6* mRNA expression and found that there was a significant downregulation of *Hdac1*, *2*, and *3*, likely contributing to its potent inhibition on class I HDACs, but no effect on *Hdac4* and *6* mRNA expression, (Supplementary Fig. 7A-E). No effect on *Tardbp* and *Ppia* mRNAs expression was also found (Supplementary Fig. 7F-G).

**FIGURE 4.**
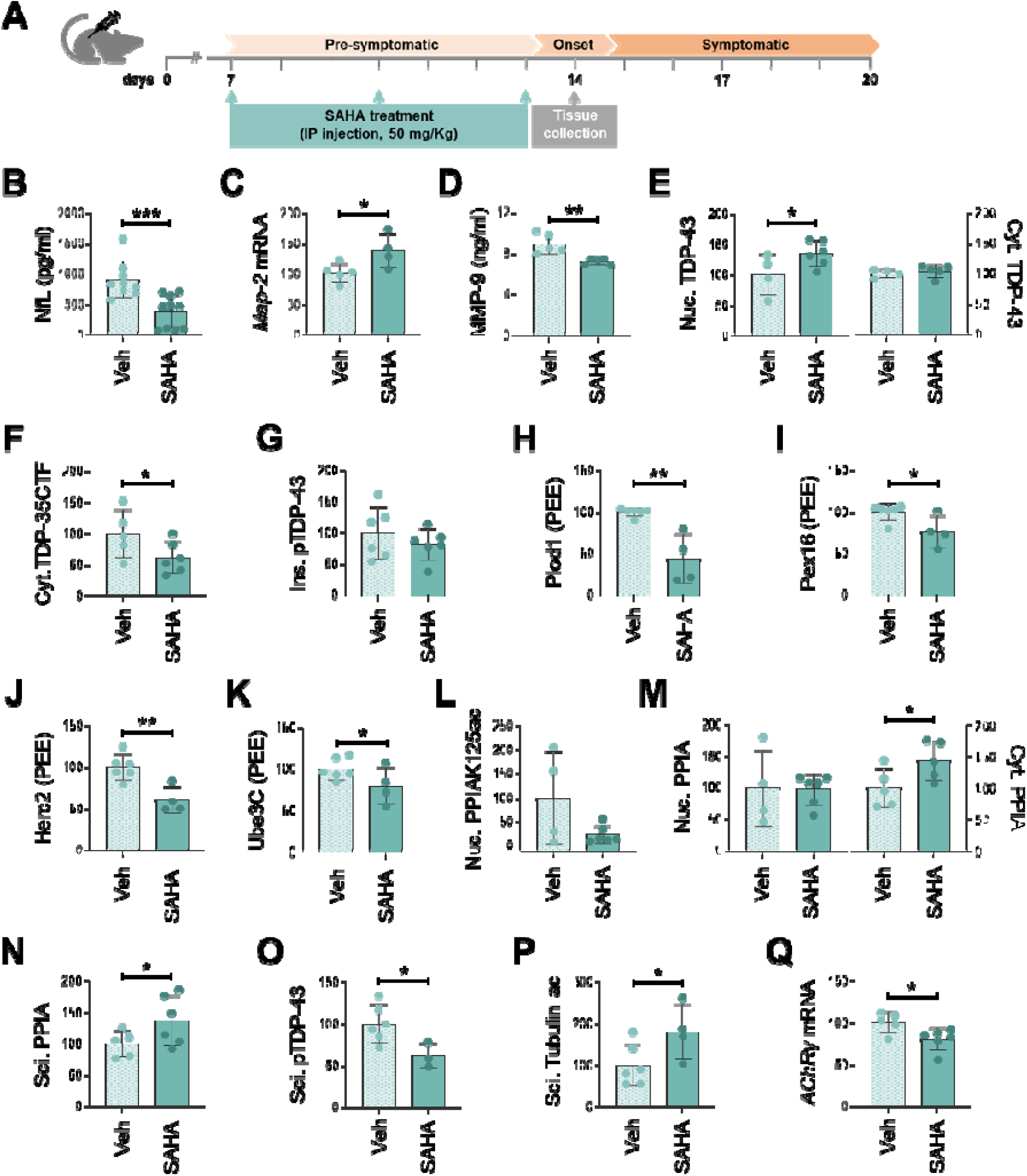
SAHA reduces neurodegeneration, TDP-43 pathology and peripheral nerve pathological markers in Thy1-hTDP-43 mouse at an early disease stage. **(A)** Scheme of SAHA treatment in homozygous Thy1-hTDP-43 mice. Mice were treated intraperitoneally with 50 mg/kg of SAHA from 7 days of age, every three days and sacrificed at 14 days for further analysis**. (B,D)** NfL **(B)** and MMP-9 **(D)** plasma levels in Thy1-hTDP-43 mice treated (SAHA; **B** n=10; **D** n=5) or not (Veh; **B** n=9; **D** n=5) with SAHA. **(C)** RT-PCR for Map-2 in spinal cord of Thy1-hTDP-43 mice treated (n=4) or not (n=5) with SAHA. Data are normalized to β-actin and reported as relative mRNA expression. **(E-G)** Analysis of nuclear (Nuc.) and cytoplasmic (Cyt.) TDP-43 **(E),** cytoplasmic TDP-35CTF **(F)** and insoluble pTDP-43 **(G)** in spinal cord of Thy1-hTDP-43 mice treated (**E** Nuc.**,F,G** n=6, **E** Cyt. n=5) or not (**E** n=4, **F** n=5, **G** Cyt. n=6) with SAHA. Data indicates dot blot **(E,G)** and WB **(F)** (WB imagines, Supplementary Fig. 6E and 9B) immunoreactivity normalized to total protein loading. **(H-K)** Semi quantitative PCR for Plod1 **(H)**, Pex16 **(I)**, Herc2 **(J)** and Ube3C (**K**) skiptic exons in spinal cord of Thy1-hTDP-43 mice treated (**H-K** n=4) or not (**H-K** n=6) with SAHA. Data are expressed as PEE. **(L-M)** Analysis of nuclear PPIAK125ac **(L)**, nuclear and cytoplasmic PPIA **(M)** in spinal cord of Thy1-hTDP-43 mice treated (**L,M** Nuc. n=6; **M** Cyt. n=5) or not (**L,M** Nuc. n=4; **M** Cyt. n=5) with SAHA. Data indicates CEI **(L)** or dot blot **(M)** immunoreactivity normalized to total protein loading **(L-M)** and to PPIA levels **(L)**. **(N-P)** Analysis of PPIA **(N)**, pTDP-43 **(O)** and acetyl tubulin (Tubulin ac) **(P)** in sciatic nerve (Sci.) of Thy1-hTDP-43 mice treated (**N** n=6; **O** n=3; **P** n=4) or not (**N** n=5; **O,P** n=6) with SAHA. Data indicates dot blot immunoreactivity normalized to total protein loading **(N-P)** and to tubulin **(P)** levels. **(Q)** RT-PCR for AChR γ-subunit in gastrocnemius muscle of Thy1-hTDP-43 mice treated (n=6) or not (n=5) with SAHA. Data are normalized to β-actin and reported as relative mRNA expression. **(B-Q)** Data are mean ± SD and are expressed as percentage of vehicle-treated mice (**C,E-Q**). *p ≤ 0.5, **p < 0.01, ***p < 0.001 by Mann Whitney test (**B,H,I**) or unpaired t test (**C-F,J,K,M-Q**).

In the second study, we treated mice with the same schedule up to 17 days of age (n=16 vehicle, n=15 SAHA) and evaluated the effect of SAHA on the same outcome measures (Fig. 5A). In this group of mice, the body weight was monitored daily confirming the delayed onset of the disease (Fig. 5B) underlined by the low level of nuclear PPIAK125ac at 14 days of age (Fig. 4L), similarly to the presymptomatic stage (Fig. 3L). In particular, ponderal growth trajectories were significantly different from 13 days of age, when vehicle-treated mice increased weight at a slower rate than the SAHA-treated mice (Fig. 5B). There was a reduced NfL concentration in plasma (Fig. 5C), however the effect on most of the other biomarkers was lost with the longer treatment (Fig. 5D-Q). There was even a tendency to an increased TDP-43 proteinopathy, no decrease of TDP-43 in the nucleus, but a significant accumulation of TDP-43 in the cytoplasm (Fig. 5F), a tendency to an increase in pTDP-43 aggregation (Fig. 5H) and no improvement in the TDP-43 splicing function (Fig. 5I-K).

**FIGURE 5.**
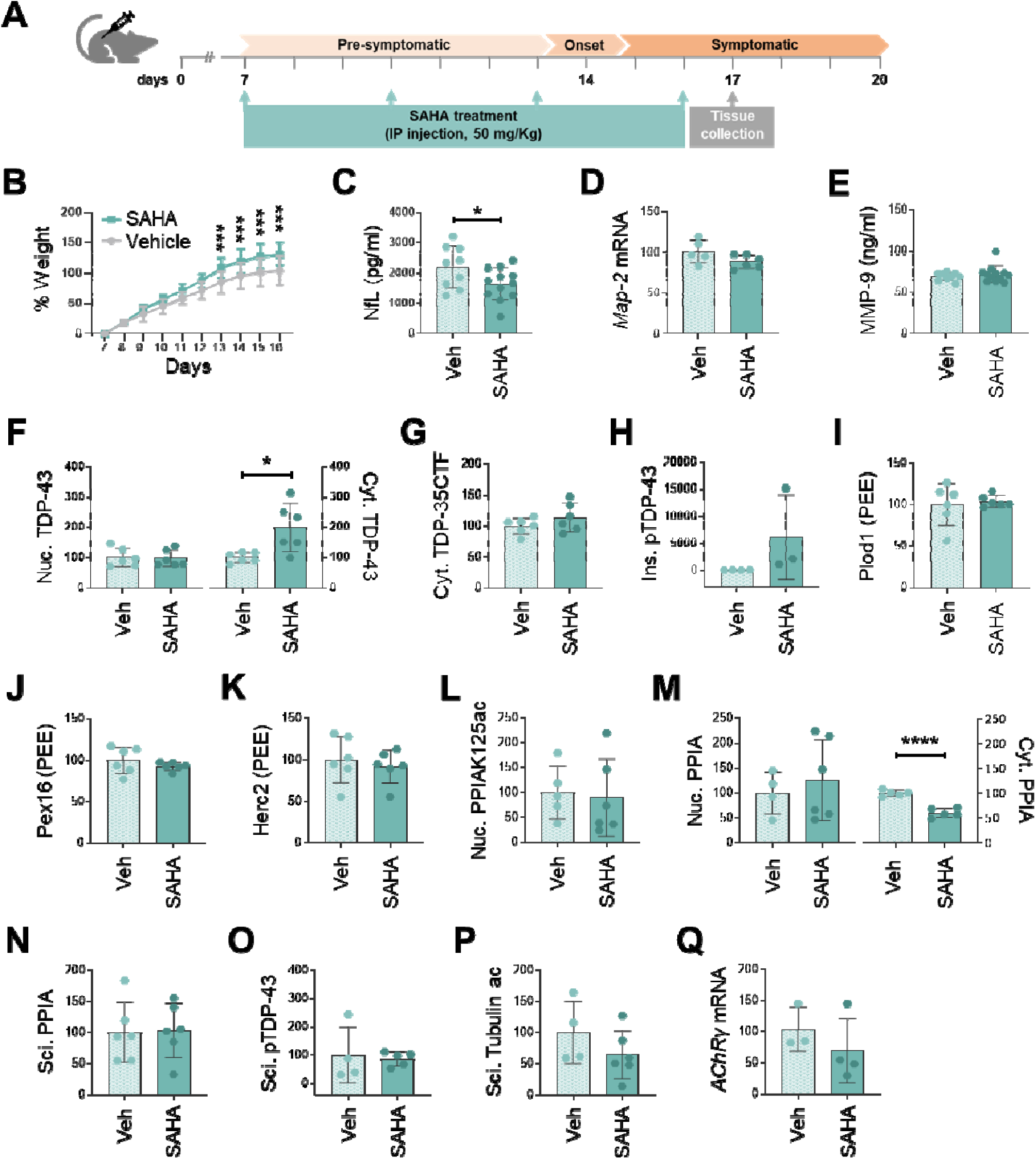
Over time, SAHA maintains low NfL levels, but the effect on most of the other biomarkers is mitigated. (**A**) Scheme of SAHA treatment in homozygous Thy1-hTDP-43 mice. Mice were treated intraperitoneally with 50 mg/kg of SAHA from 7 days of age, every three days and sacrificed at 17 days for further analysis. (**B**) Body weight curve of Thy1-hTDP-43 mice treated (n=12) or not (Veh; n=13) with SAHA. Data (mean ± SD) are expressed as percentage of body weight from 7 to 16 days of age relative to the starting body weight at 7 days of age; ***p < 0.001 vehicle versus SAHA by two-way ANOVA for repeated measures followed by Sidak’s post hoc test. (**C,E**) NfL (**C**) and MMP-9 (**E**) plasma levels in Thy1-hTDP-43 mice treated (**C**,**E** n=12) or not (**C,E** n=9) with SAHA. (**D**) RT-PCR for Map-2 in spinal cord of Thy1-hTDP-43 mice treated (n=6) or not (n=5) with SAHA. Data are normalized to β-actin and reported as relative mRNA expression. (**F-H**) Analysis of nuclear (Nuc.) and cytoplasmic (Cyt.) TDP-43 (**F**), cytoplasmic TDP-35CTF (**G**) (WB imagine, Supplementary Fig. 9C) and insoluble (Ins.) pTDP-43 (**H**) in spinal cord of Thy1-hTDP-43 mice treated (**F,G** n=6; **H** n=3) or not (**F,G** n=6; **H** n=4) with SAHA. Data indicates dot blot (**F,H**) and WB (**G**) immunoreactivity normalized to total protein loading. (**I-K**) Semi quantitative PCR for Plod1 (**I**), Pex16 (**J**) and Herc2 (**K**) skiptic exons in spinal cord of Thy1-hTDP-43 mice treated (**I-K** n=6) or not (**I-K** n=6) with SAHA. Data are expressed as PEE. (**L-M**) Analysis of nuclear PPIAK125ac (**L**), nuclear and cytoplasmic PPIA (**M**) in spinal cord of Thy1-hTDP-43 mice treated (**L,M** Nuc. n=6; **M** Cyt. n=5) or not (**L,M** Cyt. n=5; **M** Nuc. n=4) with SAHA. Data indicates CEI (**L**) and dot blot (**M**) immunoreactivity normalized to total protein loading (**L-M**) and to PPIA levels (**L**). (**N-P**) Analysis of PPIA (**N**), pTDP-43 (**O**) and acetyl tubulin (Tubulin ac) (**P**) in sciatic nerve (Sci.) of Thy1-hTDP-43 mice treated (**N,P** n=6; **O** n=5) or not (**N** n=6; **O** n=4; **P** n=4) with SAHA. Data indicates dot blot immunoreactivity normalized to total protein loading (**N-P**) and to tubulin levels (**P**). (**Q**) RT-PCR for AChR γ-subunit in gastrocnemius muscle of Thy1-hTDP-43 mice treated (n=4) or not (n=3) with SAHA. Data are normalized to β-actin and reported as relative mRNA expression. (**C-Q**) Data are mean ± SD and are expressed as percentage of vehicle-treated mice (**D,F-Q**). *p < 0.05, ****p < 0.0001 by unpaired t test (**C,F,M**).

There was no change in the level of nuclear PPIAK125ac (Fig. 5L), a decrease of PPIA in the cytoplasm with only a tendency to increase in the nucleus (Fig. 5M), likely indicating its extracellular secretion in response to the persistent cellular stress *(Pasetto* et al. *2017)*. Moreover, there was a decrease in the mRNA expression of *Hd*ac4 and *Hdac6*, not detected in the first shorter study (Supplementary Fig. 7H-L). HDAC4 plays an important role in protecting the neuromuscular function in ALS *(Pigna* et al. *2019; Boutillier* et al. *2019)*, while HDAC6 is involved in the clearance of misfolded proteins (*Lee* et al. 2010; *Kawaguchi* et al. *2003*). This may suggest that longer SAHA treatment could not be effective also because of an accumulation of a number of undesirable side effects.

*In vitro* studies in ALS models have reported that SAHA may act as an HSP co-inducer and that this effect is potentiated by arimoclomol, enhancing HSP70 expression in combination (Kuta *et al*. 2020; Fernández Comaduran *et al*. 2024) We therefore tested if the combination of the two drugs were more effective in Thy1-hTDP-43 mice in counteracting the 2-fold hTDP-43 overexpression and compensate for the side effects. We treated mice with SAHA+arimoclomol (n=17) or vehicle (n=15), with the same schedule up to 17 days of age and evaluated the effect of the treatment on the outcome measures (Fig. 6A). Ponderal growth trajectory data did not show significant differences between treated and vehicle-treated animals (Fig. 6B). However, the treatment increased HSP70 level in the spinal cord and respect to SAHA alone had a stronger neuroprotective effect, with a greater decrease in NfL (−25% with SAHA versus −38% with SAHA+arimoclomol), and a significant decrease of MMP-9 in plasma (Fig. 6C,D,F). Although TDP-43 pathology in the spinal cord did not greatly improve, no worsening was observed (Fig. 6G-I), as seen with SAHA alone (Fig. 5F-H), and an improvement in the TDP-43 splicing function was detected, with a significant decrease in the exclusion of the skiptic exons for Ube3C and Slc6a6 (Fig. 6J-K). No significant accumulation of cytoplasmic TDP-43 compared to vehicle-treated mice (Fig. 6G versus Fig. 5F) and no significant decrease in cytoplasmic PPIA were observed (Fig. 6M versus Fig. 5M). Moreover, SAHA+arimoclomol did not affect *Hdac4* and *Hdac6* mRNA expression (Supplementary Fig. 7P-Q), indicating that possible negative effects from *Hdac4* and *Hdac6* downregulation were rescued. Interestingly, a great improvement was seen in periphery, regardless of no change in PPIA (Fig. 6N). Mice treated with the combination showed a substantial increase in acetylated tubulin, an equally important decrease in pTDP-43 in the sciatic nerve and a significant decrease in the *AChR* γ-subunit expression in the gastrocnemius muscle (Fig. 6O-Q). Arimoclomol alone was not affecting acetylation of tubulin, suggesting a possible synergistic effect of the two drugs on HDAC6 inhibition (Supplementary Fig. 8B). Moreover, arimoclomol did not reduce pTDP-43 levels in the sciatic nerve as the combination (Supplementary Fig. 8C). Arimoclomol reduced NfL and *AChR* γ-subunit expression, although not to the same extent as the combination (Supplementary Fig. 8D-E). Overall, arimoclomol potentiated the positive effects of SAHA in periphery and mitigated its side effects.

**FIGURE 6.**
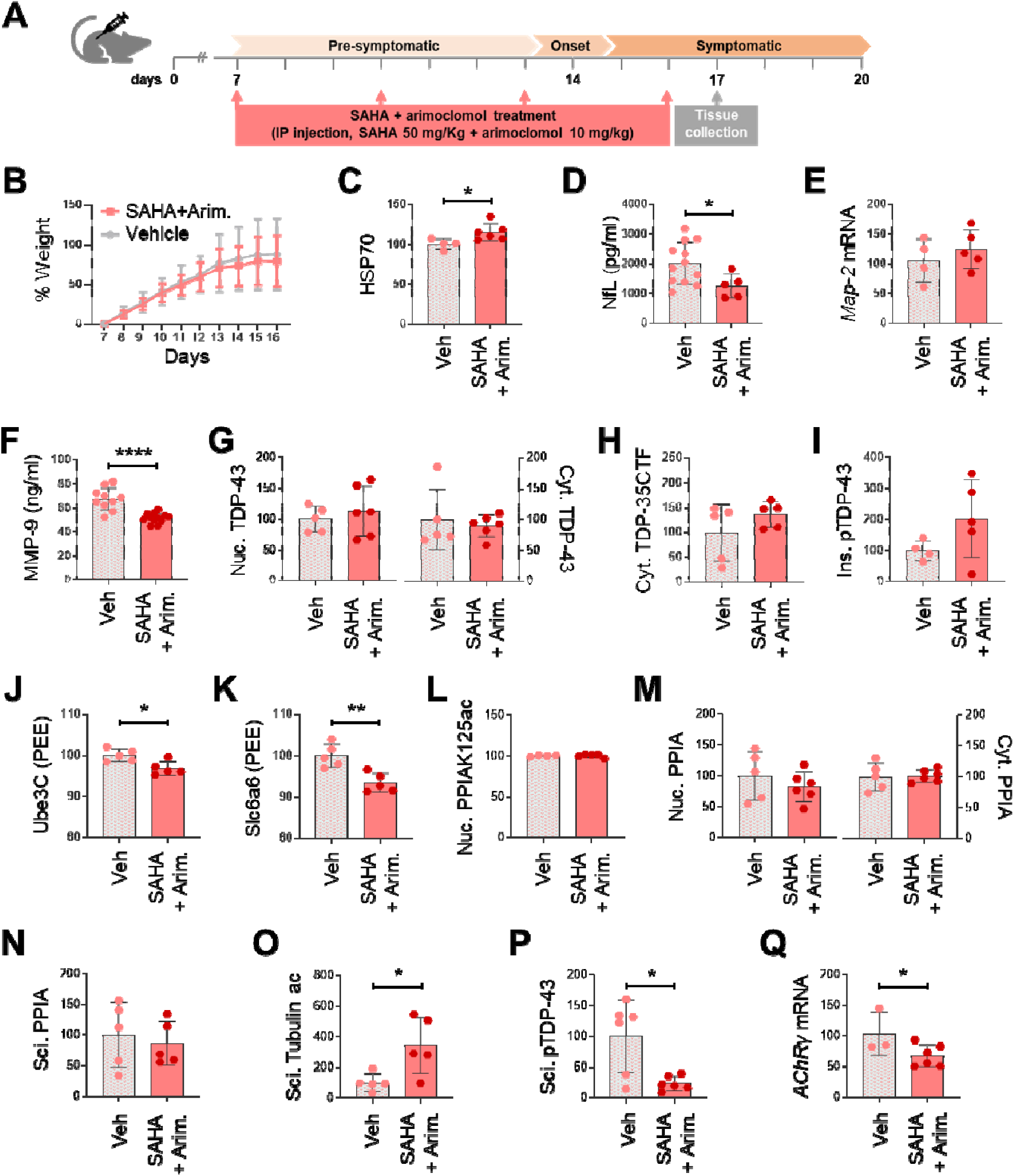
SAHA+Arimoclomol enhances the positive effect on markers of neuromuscular pathology. (**A**) Scheme of SAHA+Arimoclomol (SAHA+Arim.) treatment combination in homozygous Thy1-hTDP-43 mice. Mice were treated intraperitoneally with 50 mg/kg of SAHA and 10 mg/kg of arimoclomol, from 7 days of age, every three days and sacrificed at 17 days for further analysis. (**B**) Body weight curve of Thy1-hTDP-43 mice treated (n=12) or not (Veh; n=10) with SAHA+Arim. Data (mean ± SD) are expressed as percentage of body weight from 7 to 16 days of age relative to the starting body weight at 7 days of age; two-way ANOVA for repeated measures followed by Sidak’s post hoc test. (**C**) Level of HSP70 in spinal cord of Thy1-hTDP-43 mice treated (n=6) or not (Veh; n=4) with SAHA+Arim (WB imagine, Supplementary Fig. 9D). Data indicates WB immunoreactivity normalized to total protein loading. (**D,F**) NfL (**D**) and MMP-9 (**F**) plasma levels in Thy1-hTDP-43 mice treated (**D** n=5; **F** n=12) or not (**D** n=12; **F** n=4) with SAHA+Arim. (**E**) RT-PCR for Map-2 in spinal cord of Thy1-hTDP-43 mice treated (n=5) or not (n=4) with SAHA+Arim. Data are normalized to β-actin and reported as relative mRNA expression. (**G-I**) Analysis of nuclear (Nuc.) and cytoplasmic (Cyt.) TDP-43 (**G**), cytoplasmic (Cyt.) TDP-35CTF (**H**) (WB imagine, Supplementary Fig. 9E) and insoluble (Ins.) pTDP-43 (**I**) in spinal cord of Thy1-hTDP-43 mice treated (**G** n=6; **H,I** n=5) or not (**G,H** n=5; **I** n=4) with SAHA+Arim. Data indicates dot blot (**G,I**) and WB (**H**) immunoreactivity normalized to total protein loading. (**J-K**) Semi quantitative PCR for Ube3C (**J**) and Slc6a6 (**K**) skiptic exons in spinal cord of Thy1-hTDP-43 mice treated (**J-K** n=5) or not (**J-K** n=5) with SAHA+Arim. Data are expressed as PEE. (**L-M**) Analysis of nuclear (Nuc.) PPIAK125ac (**L**), nuclear (Nuc.) and cytoplasmic (Cyt.) PPIA (**M**) in spinal cord of Thy1-hTDP-43 mice treated (**L** n=5; **M** n=6) or not (**L** n=4; **M** n=5) with SAHA+Arim. Data indicates CEI (**L**) and dot blot (**M**)immunoreactivity normalized to total protein (**L-M**) and to PPIA levels (**L**). (**N-P**) Analysis of PPIA (**N**), acetyl tubulin (Tubulin ac) (**O**) and pTDP-43 (**P**) in sciatic nerve (Sci.) of Thy1-hTDP-43 mice treated (**N,O** n=5; **P** n=6) or not (**N,O** n=5; **P** n=6) with SAHA+Arim. Data indicates dot blot immunoreactivity normalized to total protein loading (**N-P**) and to tubulin levels (**O**). (**Q**) RT-PCR for AChR γ-subunit in gastrocnemius muscle of Thy1-hTDP-43 mice treated (n=6) or not (n=3) with SAHA+Arim. Data are normalized to β-actin and reported as relative mRNA expression. (**C-Q**) Data are mean ± SD and are expressed as percentage of vehicle-treated mice (**C,E,G-Q**). *p < 0.5, **p < 0.01, ****p < 0.0001 by unpaired t test (**C,D,F,J,K,O-Q**).

## 4 Discussion

HDACis are valuable tools to explore acetylation homeostasis and hold potential for treating neurodegeneration, but they have been scarcely investigated in the context of TDP-43 pathology. Several of them are in advanced stages of development or have been approved for cancer therapy, such as SAHA, therefore they might be suitable candidates for repurposing. However, it is important to emphasize that HDACis are potent drugs that induce extensive transcriptional changes and should therefore be carefully studied in the specific disease models throughout the progression of the disease to derive the net effect of their multiple actions. Our interest in HDACis stems from the observation that Lys-acetylation is involved in the interaction of TDP-43 with PPIA, which has a key role in regulating TDP-43 localization and function (Lauranzano *et al*. 2015). In this work we found that HDAC1 can deacetylate PPIA and that, inhibiting this pathway by SAHA, TDP-43 proteinopathy is reduced in a cellular model of chronic nutrient starvation and in PBMCs of ALS patients. SAHA also delayed the onset of TDP-43 pathology in an aggressive mouse model of TDP-43 proteinopathy, but the effect was temporary. When in combination with arimoclomol a mitigation of the neurodegeneration was maintained over time. Moreover, there was a great reduction of the pathological markers in the sciatic nerve, including pTDP-43, and markers of axonal stability and muscle denervation.

HDAC inhibition has been largely investigated *in vitro* and in preclinical trials using several mouse models of ALS (Klingl *et al*. 2021), but not in mouse models of TDP-43 proteinopathy and in combination with arimoclomol. While we were submitting our paper, a study was published reporting the effect of an HDAC6 inhibitor, EKZ-438, in the rNLS8 TDP-43ΔNLS mouse, showing improved TDP-43 proteostasis specifically in the brain (James *et al*. 2025). Our aim was to investigate how acetylation homeostasis influences the early events driving TDP-43 pathology, which likely result in impaired nucleocytoplasmic transport and loss of nuclear TDP-43, disruptions associated with widespread dysfunctions in RNA metabolism. In our *in vitro* experiments, SAHA turned out to be effective in inhibiting accumulation of TDP-43 and TDP-35CTF in the cytoplasm. SAHA is considered a broad-spectrum inhibitor since can strongly inhibit class I enzymes, such as HDAC1 and HDAC3, but also class II enzymes, such as HDAC6, although less efficiently (Witter *et al*. 2008; Padige *et al*. 2015; Jones *et al*. 2008; Huber *et al*. 2011). An interactomic study found that PPIA interacts with HDAC1 (Maneix *et al*. 2024) and our siRNA experiments indicate that PPIA is deacetylated by HDAC1. Moreover, HDAC1 reduction and inhibition was protective against mutant TDP-43 toxicity (Sanna *et al*. 2020). This suggests a primarily involvement of an HDAC1-dependent mechanism, but we cannot exclude the contribution of the inhibition of other HDACs, especially HDAC6. Therefore, our results are not in disagreement with reported data *in vivo* and *in vitro* with HDAC6 inhibitors (James *et al*. 2025; Fazal *et al*. 2021; Guo *et al*. 2017; Rossaert *et al*. 2019). The effect of SAHA on TDP-43 localization was also confirmed in ALS patient PBMCs. These cells have shown to display cytoplasmic accumulation of TDP-43 in a previous study (De Marco *et al*. 2011). In a later study, we have also shown that, in PBMCs from a relative large cohort of ALS patients, TDP-43 was entrapped in supramolecular assemblies with hnRNPA2B1 and PPIA (Luotti *et al*. 2020). Altogether, these findings suggest that PBMCs may be useful to test the effect of therapeutic approaches targeting TDP-43 proteinopathy in patients and possible monitor clinical trials.

To test the effect of SAHA on the onset of TDP-43 proteinopathy *in vivo,* we chose a model with marked pathology induced in neurons by hTDP-43 overexpression, in absence of a specific mutation. The mice show neuromuscular junction (NMJ) pathology, denervation of distal muscles and motor neuron loss, as observed in ALS patients (Wils *et al*. 2010; Alhindi *et al*. 2023). Moreover, the timing of the onset of the proteinopathy is well defined and, because of the short life span, preclinical studies with multiple treatment schedules and drug combinations are feasible. However, due to the neonatal onset and severity of the disease, behavioral studies on disease progression are limited. Therefore, we performed multiple molecular evaluations at different stages of the disease to identify surrogate markers. Interestingly, we could monitor TDP-43 impairment in splicing function with the progression of the disease and in response to SAHA treatment. In particular, we evaluated known skiptic exons, i.e. excision of otherwise normally conserved exons, and cryptic exons, i.e. decreased excision of non-conserved cryptic exons, that are the result of TDP-43 dysfunction (Fratta *et al*. 2018; Ling *et al*. 2015). We could demonstrate that in conditions of transgenic overexpression of the hTDP-43 protein, there is no loss of the TDP-43 cryptic exon excision function, while increased exon skipping can be seen already at an early stage of the disease and seems to be a good proxy of nuclear TDP-43 levels. It has been hypothesized that exclusion of highly conserved skiptic exons in an early phase may affect key cellular pathways such as protein degradation (Ube3C, Herc2, Pacrgl) and inflammatory response (Ankrd42) that in turn could drive TDP-43 aggregation, nuclear depletion and loss of TDP-43 function (Fratta *et al*. 2018; Rouaux *et al*. 2018). In a second phase, this may lead to increasing occurrence of cryptic exons, which possibly are not observable in a mouse model with such a fast disease progression.

In PBMCs of ALS patients we could correlate reduced TDP-43 proteinopathy with increased nuclear PPIAK125ac due to lysine deacetylation inhibition with SAHA. In the mice this effect could be also associated with an improved splicing function of TDP-43 only in the short-term study. Instead, with the longer treatment the positive effects were mitigated, possibly because SAHA has an impact especially on TDP-43 trafficking at the onset of TDP-43 proteinopathy. However, over-time, its effect is insufficient to counteract the 2-fold overexpression of hTDP-43 in the neurons of Thy1-hTDP-43 mice. Moreover, with the longer treatment, the broad-spectrum HDAC inhibition by SAHA may induce side effects specifically in the nervous system. We found downregulation of HDAC4 and HDAC6. Other studies reported an inhibitory effect of SAHA on HDAC4 (Ho *et al*. 2020), which has an essential role in synaptic plasticity and memory formation (Kim *et al*. 2012), and in compensatory muscle reinnervation during ALS (Pigna *et al*. 2019). SAHA inhibits also HDAC6, although not greatly (IC_50_ >30 nM) (Witter *et al*. 2008; Padige *et al*. 2015; Jones *et al*. 2008; Huber *et al*. 2011). Its inhibition favors axonal transport (Fazal *et al*. 2021) and axonal outgrowth (Kalinski *et al*. 2019), but at the same time may increase TDP-43 acetylation, a determinant of TDP-43 aggregate formation (Cohen *et al*. 2015). The latest aspect is particularly critical in a condition of high susceptibility to TDP-43 proteinopathy and TDP-43 overexpression.

To overcome overexpression and consequent aggregation of TDP-43 in the mouse model, we tested the SAHA+arimoclomol combination. Arimoclomol is a small drug that prolongs the activation and the DNA binding of the HSF-1 transcription factor, upregulating the molecular chaperone family of HSPs (i.e. Hsp70, Hsp90). Although arimoclomol was effective at ameliorating disease phenotypes in multiple preclinical models of motor neuron disease (Kieran *et al*. 2004; Kalmar *et al*. 2008; Ahmed *et al*. 2023), it failed to modify the progression of the disease in a phase 3 clinical trial for ALS (Benatar *et al*. 2024). The combination SAHA+arimoclomol is known to have a synergistic effect in enhancing stress-induced Hsp70 expression in spinal cord-dorsal root ganglion cultures in response to proteotoxic stress (Kuta *et al*. 2020). Moreover, RGFP963, another class I HDACi, in combination with arimoclomol significantly increased axonal transport in cultures expressing mutant SOD1 (Fernández Comaduran *et al*. 2024). The mechanisms by which HDAC inhibition and HSP co-inducers interact and enhance HSP response is complex and not fully elucidated. While HDAC inhibitors have shown HSP-inducing properties, arimoclomol did not inhibit class I HDACs using an *in vitro* assay (Kuta *et al*. 2020). Moreover, arimoclomol can exert neuroprotective effects without altering HSP expression, suggesting the involvement of alternative mechanisms (Fernández Comaduran *et al*. 2024; Pelaez *et al*. 2024). Thus, the mechanism by which arimoclomol potentiated SAHA in the present study requires further investigations. In this work, we provided evidence that the combination, tested for the first time *in vivo,* boosted neuroaxonal and neuromuscular protection, without full recovery from TDP-43 proteinopathy and only a partial rescue of the splicing function. The combination SAHA+arimoclomol was more effective in decreasing markers of neurodegeneration and neuroinflammation (NfL, MMP-9), but especially greatly reduced markers of neuromuscular pathology in sciatic nerve and gastrocnemius muscle: tubulin deacetylation, accumulation of pTDP-43 and AChR γ-subunit re-expression. Acetylation of α-tubulin at Lys-40 is a marker of microtubule stability and promotes the recruitment of the molecular motors dynein and kinesin-1 to microtubules, promoting anterograde and retrograde transport (Dompierre *et al*. 2007). In fact, increasing tubulin acetylation through HDAC6 inhibition has improved axonal transport in models of ALS and Charcot-Marie-Tooth Disease (d’Ydewalle *et al*. 2012; Taes *et al*. 2013). This in turn may favor axonal transport of TDP-43 mRNP granules and facilitate delivery of TDP-43 mRNA targets to distal neuronal compartments (Alami *et al*. 2014), thus explaining the reduction of pTDP-43 accumulation in the sciatic nerve. Previous studies have shown that NMJ innervation of muscles was better preserved by either HDAC6 inhibition or arimoclomol treatment individually (Kieran *et al*. 2004; Taes *et al*. 2013). In the fetal AChR isoform of skeletal muscle endplates, the γ-subunit is downregulated after birth and replaced by the ε-subunit, except in cases of denervation, where the fetal form is re-expressed in the muscle. Indeed, downregulation of the γ-subunit has been detected in ALS mice following treatments that preserve innervation and muscle mass in the hindlimbs (Trolese *et al*. 2022; Fabbrizio *et al*. 2022). Here, in the longer treatment, we observed only a tendency to decrease in AChR γ-subunit expression following treatment with either SAHA or arimoclomol individually, while the effect was significant when the two were used in combination. This result is particularly notable given that Thy1-hTDP-43 mice exhibit pronounced NMJ denervation in hindlimb muscles and NMJ pathology is restricted to motor nerve terminals (Alhindi *et al*. 2023).

There is an increasing interest in testing drug cocktails, especially with repurposed drug candidates, because of the multifactorial nature of ALS. Our results highlight the importance of *in vivo* investigations for addressing the complexity of drug mechanisms, especially in combination therapies, to reveal unpredictable negative or positive effects of drug-drug interaction in specific tissues/organs and stages of the disease, which may be overlooked in cell culture studies. Although the Thy1-hTDP-43 mouse is useful to decipher early molecular mechanisms at the basis of TDP-43 proteinopathy and NMJ denervation, has an excessively aggressive disease and is limited in evaluating the effects of drugs with the progression of the disease. Experiments with a mouse model with a slower and adult-onset disease are warranted.

Regarding the role of PPIA, these studies further suggest that TDP-43 homeostasis is critically influenced by PPIA. Increased acetylation on K125 is associated with a reverse TDP-43 localization in a cellular model of TDP-43 proteinopathy and in PBMCs of ALS patients, where TDP-43 likely undergoes liquid demixing, but does not irreversibly aggregate. Moreover, a reduction of insoluble pTDP-43 was also observed *in vitro* by transfecting the PPIA acetylation-mimetic mutant. From the studies in mice, it appeared that PPIA125Kac is involved in an early response that neurons adopt to counteract TDP-43 altered homeostasis. It is possible that PPIA and its acetylated form accumulate with TDP-43 in membrane-less compartments in response to stress, as reported in other paradigms (Yu *et al*. 2021; Lu *et al*. 2022; Curdy *et al*. 2023; Xiang *et al*. 2015), and that persistent stress may lead to irreversible protein aggregation, stress-induced PPIA secretion and finally loss of a key intracellular protective factor. In fact, we previously demonstrated high PPIA in CSF and plasma of patients and animal models, low intracellular soluble PPIA and high detergent-insoluble protein in CNS tissues of ALS mouse models and ALS/FTD patients (Pasetto *et al*. 2017; Luotti *et al*. 2020; Basso *et al*. 2009; Pasetto *et al*. 2021).

However, this work does not allow us to definitively establish a causal relationship between PPIA acetylation and the improvement of TDP-43 homeostasis. Increased PPIA acetylation was achieved using a broad-spectrum HDAC inhibitor, which may exert multiple effects through transcriptional modulation. Experiments aimed at testing an acetylated PPIA mimetic in ALS animal models are ongoing in the laboratory to address this question.In conclusion, the present study suggests that HDAC inhibitioncould be beneficial for restoring TDP-43 localization and function, through multiple mechanisms, including modulation of PPIA acetylation. SAHA, a broad-range HDAC inhibitor, reduced TDP-43 proteinopathy in a severe and fast progressing TDP-43 overexpressing mouse model, although transiently. Arimoclomol in combination with SAHA, mitigated side effects and showed a synergistic action in periphery, significantly reducing markers of peripheral nerve pathology and muscle denervation. Therefore, it is worthwhile to further explore lysine deacetylation inhibition and arimoclomol in combination, as a potential therapeutic approach for patients.

## Data availability

Data that support the findings of this study are available within the paper and Supplementary material. Raw data have been deposited in the Zenodo public repository (https://zenodo.org/).

## Abbreviations

AD: Alzheimer’s disease
ALS: Amyotrophic lateral sclerosis
AChR: acetylcholine receptor
CEI: Capillary electrophoresis immunoassay
FTD: frontotemporal dementia
HDACs: histone deacetylases
HDACis: HDAC inhibitors
HSP: heat shock protein
MMP-9: matrix metalloproteinase-9
NfL: neurofilament light chain
PBMCs: peripheral blood mononuclear cells
PPIA: Peptidyl-prolyl cis-trans isomerase A
PPIAK125ac: K125 acetylated PPIA
SAHA: suberoylanilide hydroxamic acid
TDP-43: TAR DNA-binding protein 43 kDa
TDP-35CTF: C-terminal fragment of 35 kDa
WB: Western Blot

## Acknowledgements

We would like to express our sincere gratitude to the patients who generously agreed to participate in this study.

## Authors’ information

Serena Scozzari and Stefano Fabrizio Columbro contributed equally to this study.

## Contributions

VB and LP were responsible for conception and design of the study. All authors contributed to data acquisition and analysis. VB and LP contributed to drafting the text and preparing the figures. All authors critically evaluated and approved the final manuscript.

## Ethics Declarations

### Ethics Approval and Consent to Participate

Subjects enrolled in this study signed a written informed consent before blood drawn. The study was supported by the Italian Ministry of Health and approved by the Ethics Committee of the Azienda Ospedaliero-Universitaria Città della Salute e della Scienza di Torino (protocol number 0062147 June 19, 2019), in accordance with the ethical standards laid down in the 1964 Declaration of Helsinki and its later amendments.

