## Supplementary Figures and Table for "Effects of lysine deacetylation inhibition alone or in combination with arimoclomol on TDP-43 proteinopathy": Supplementary Figures_2026.pdf

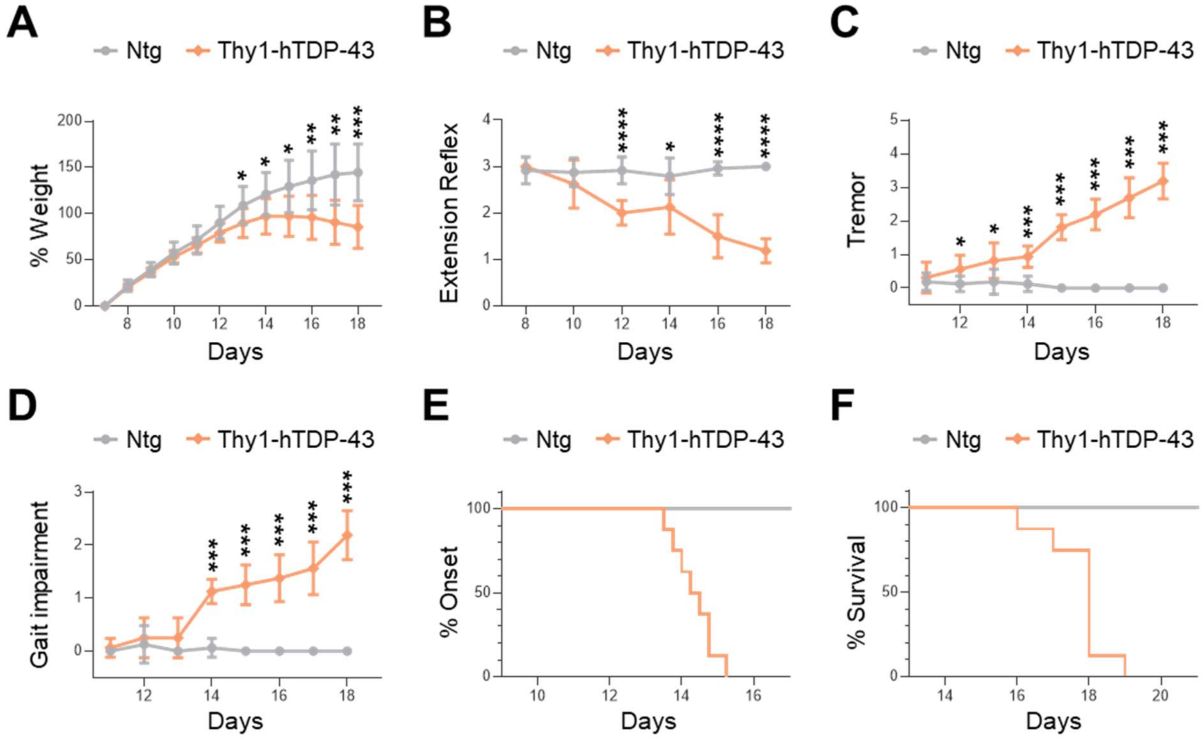


**Supplementary Figure 1. Behavioral characterization of Thy1-hTDP-43 mouse model. (A)** Body weight curve of Ntg (n=12) and Thy1-hTDP-43 mice (n=8). Data are expressed as percentage of body weight from 7 to 18 days of age normalized to the body weight at 7 days of age. **(B-D)** Evaluation of hind limb extension reflex **(B)**, tremor **(C)** and gait impairment **(D)** score for Ntg and Thy1-hTDP-43 mice during disease progression. **(A-D)** Data are mean ± SD (n=8-12 in each group). *$p \leq 0.5$, **$p < 0.01$, ***$p < 0.001$ and ****$p < 0.0001$, Ntg versus Thy1-hTDP-43 by two-way ANOVA for repeated measures followed by Tukey's post hoc test (**B**) or by multiple Mann-Whitney tests (**A,C,D**). **(E-F)** Kaplan–Meier curve for disease onset **(E)** and survival **(F)** of Ntg and Thy1-hTDP-43 mice (n=8-12 in each group). ****$p < 0.0001$, Ntg versus Thy1-hTDP-43 mice by Log-rank Mantel-Cox test (median ± SD: onset 14 ± 0.69 days; survival 18 ± 0.89 days).

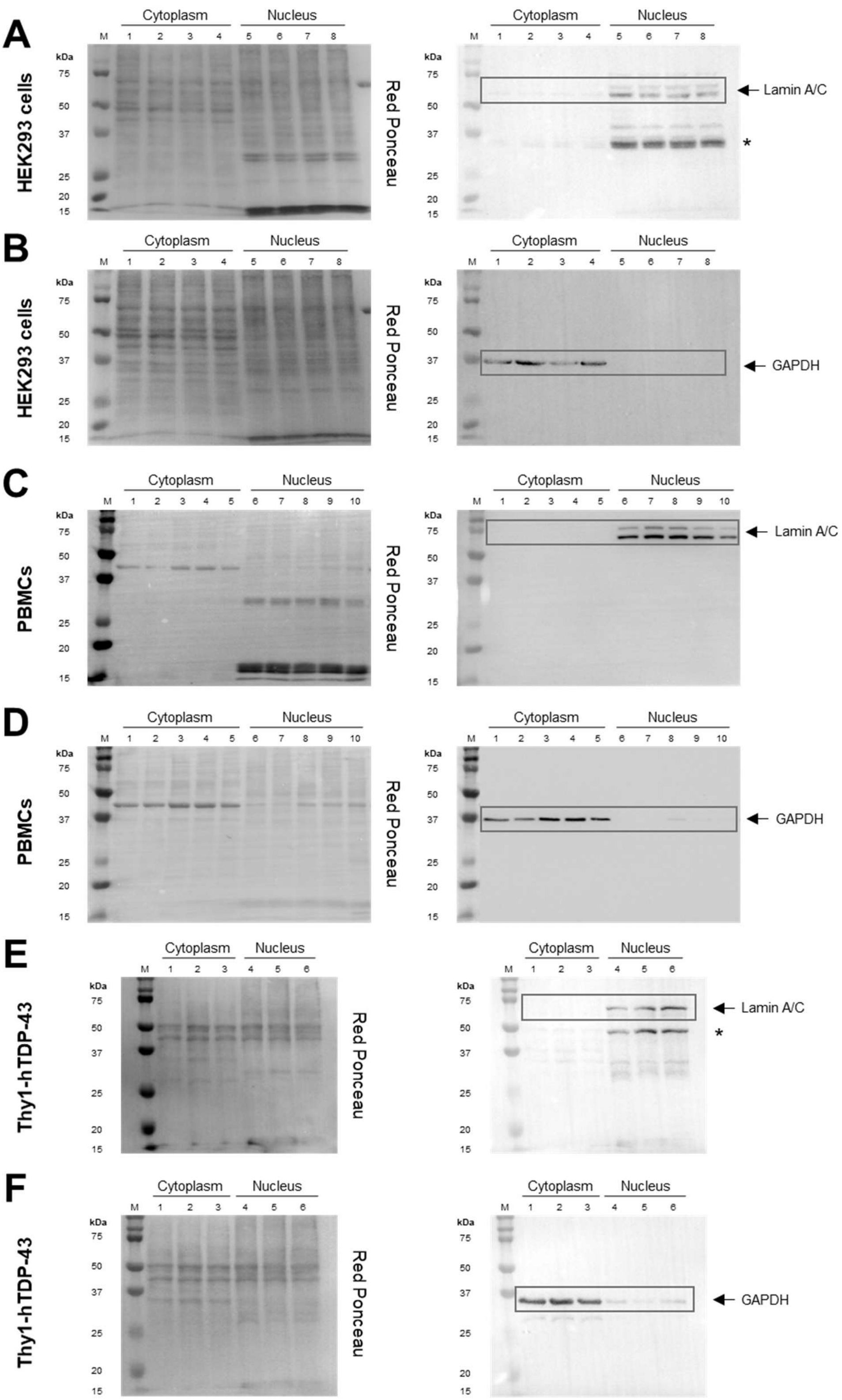


**Supplementary Figure 2. Assessment of subcellular fractionation in HEK293 cell cultures, human PBMCs and murine spinal cord. (A-F)** Representative WB after Red Ponceau staining (left) and antibody hybridization (right) for lamin A/C **(A,C,E)**, nuclear marker, or GAPDH **(B,D,F)**, cytoplasmic marker, in nuclear and cytoplasmic fractions obtained from control HEK293 cells **(A-B)**, human PBMCs of ALS patients **(C-D)** and spinal cord tissue of Thy1-hTDP-43 mice **(E-F)**. *Asterisks indicate bands that likely correspond to cleaved forms of lamin A/C, large (41-50 kDa) and small (28 kDa) fragments.

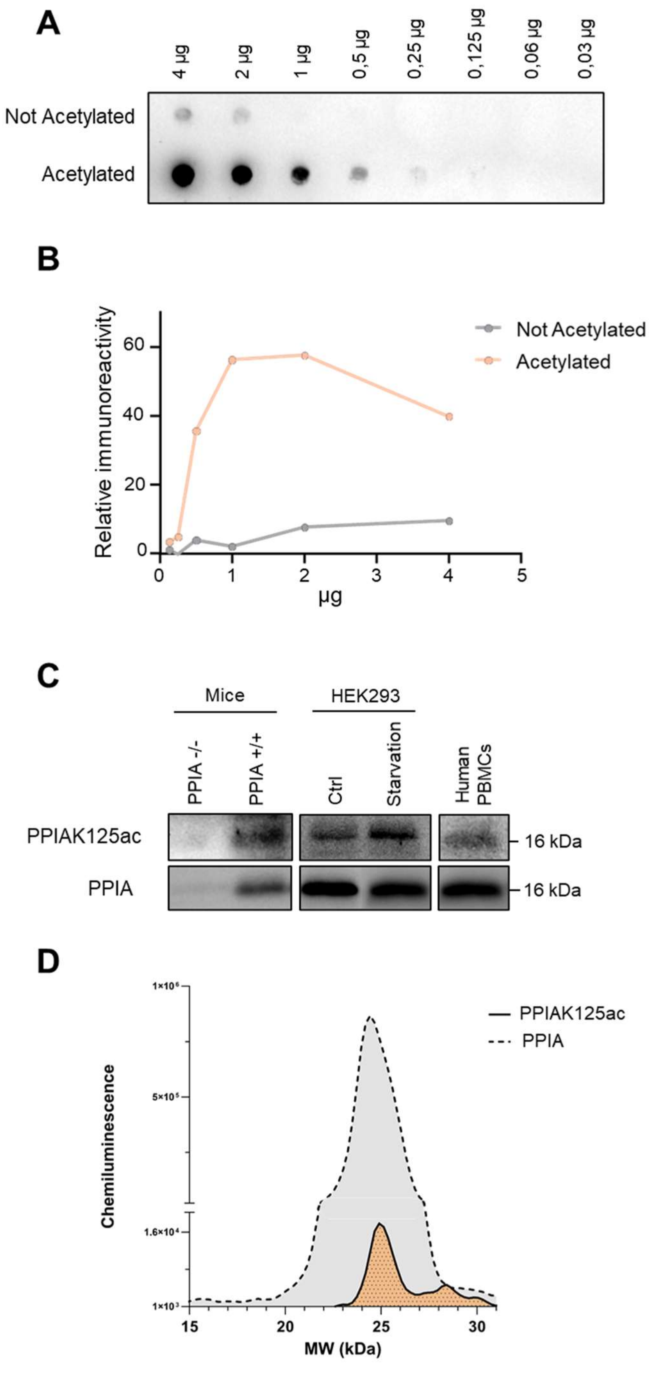


**Supplementary Figure 3. Characterisation of acetyl-PPIA antibody. (A)** Representative dot blot of different amount of the recombinant peptide WLDGKHVVFG, acetylated or not-acetylated at the K residue, after anti-PPIAK125ac antibody hybridization. **(B)** Dose-dependent relative immunoreactivity of acetylated (orange) versus not acetylated (grey) peptide analyzed at increasing amounts (0.03–4 µg) and normalized on Red Ponceau staining. **(C)** Representative WB after hybridization with anti-PPIAK125ac and PPIA antibody of spinal cord tissues from PPIA knock-out (PPIA-/-) mice and relative Ntg controls, lysate of serum-deprived HEK293 cells (Starvation) and control (Ctrl) and human PBMCs from ALS patients. **(D)** Representative electropherogram following PPIAK125ac and PPIA hybridization in the same capillary through CEI in spinal cord of Ntg mice. The peak of PPIA (grey) includes that of PPIAK125ac (orange).

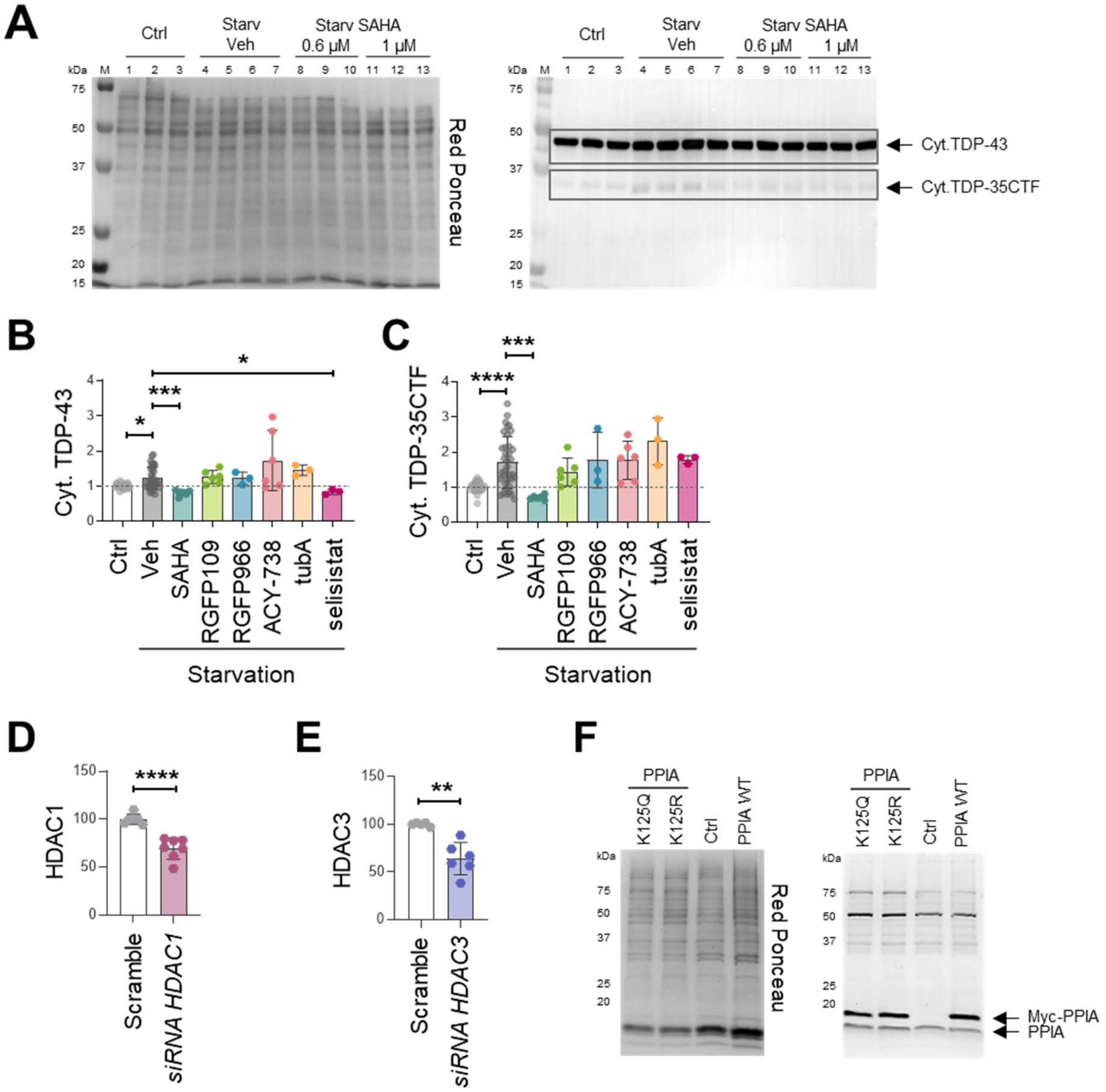


**Supplementary Figure 4. The effects of SAHA in an *in vitro* model of TDP-43 proteinopathy. (A)** Uncropped blot of representative WB shown in Figure 1A. WB after Red Ponceau staining (left) and antibody hybridization (right) for TDP-43 in cytoplasmic fraction derived from control and serum-deprived HEK293 cells (Starv), vehicle-treated (Veh) or treated with 0.6 and 1 µM SAHA. Arrows and boxes indicate the bands used for quantification. **(B-C)** Cytoplasmic (Cyt.) levels of TDP-43 (**B**) and TDP-35CTF (**C**) in control (Ctrl; n ≥ 6 independent cell culture preparations) and serum-deprived HEK293 cells (Starvation), vehicle-treated (Veh; n ≥ 6 independent cell culture preparations) or treated with 1 µM HDACi (n=3-6 independent cell culture preparations): SAHA, RGFP109, RGFP966, ACY-738, tubastatin A (tubA) and selisistat. Dashed line indicates Ctrl level. Each HDACi and Veh condition was normalized to its respective Ctrl from the corresponding independent experiment. **(D-E)** Quantification of HDAC1 **(D)** or HDAC3 **(E)** levels in total lysate of HEK293 transiently transfected with siRNA control (Scramble; n=5-6 independent cell culture preparations) and siRNA for *HDAC1* (n=7 independent cell culture preparations), or *HDAC3* (n=6 independent cell culture preparations). (**F**) Myc immunoreactivity and relative Red Ponceau staining in HEK293 cells transfected with Myc-PPIA mutants K125Q, K125R, WT and in HEK293 Ctrl cells. Arrows indicate the immunosignal of transfected Myc-PPIA, as well as the endogenous PPIA signal related to the previous hybridization with the anti-PPIA antibody. Data are expressed as percentage of scramble levels. **(B-E)** Data (mean ± SD) indicates WB **(B-C)** and CEI **(D-E)** immunoreactivity normalized to total protein loading. *p < 0.05, **p < 0.01, ***p < 0.001, ****p < 0.0001 by one-way ANOVA with Dunnett's multiple comparisons post hoc test (**C**), by Kruskal-Wallis with Dunnett's multiple comparisons post hoc test (**B**) or by unpaired t test (**D-E**).

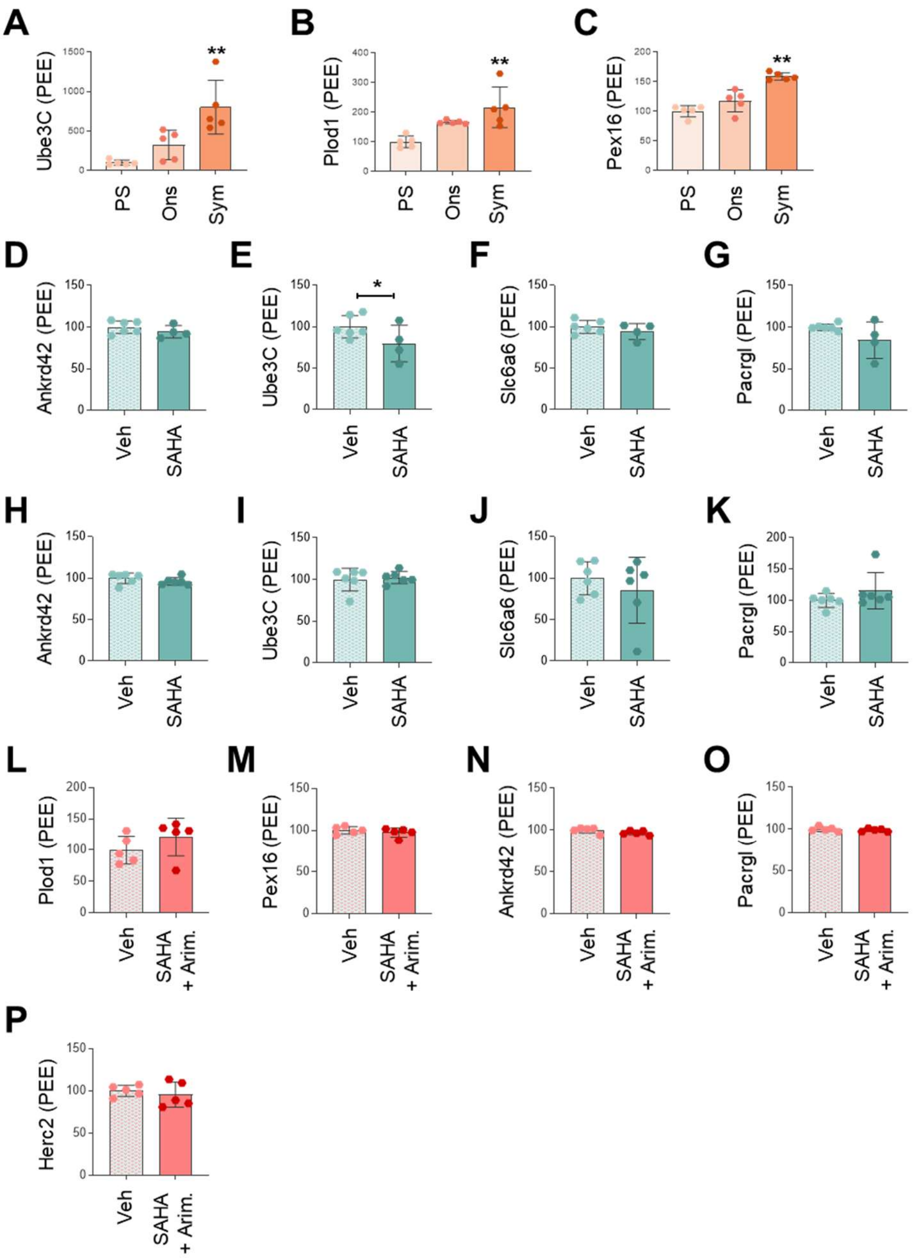


**Supplementary Figure 5. Analysis of a set of skiptic exons in the spinal cord of homozygous Thy1-hTDP-43 mice. (A-C)** Semi quantitative PCR of Ube3C **(A)**, Plod1 **(B)**, Pex16 **(C)** skiptic exons in spinal cord of Thy1-hTDP-43 at pre-symptomatic (PS, 7 days of age), onset (Ons, 14 days of age) and symptomatic (Sym, 17 days of age) stage. Data (mean ± SD; n =5 in each experimental group) are expressed as PEE. **p < 0.01 versus PS by one-way ANOVA with Tukey's multiple comparisons (**B**) or by Kruskal-Wallis with Dunn's multiple comparisons test (**A,C**). **(D-G)** Semi quantitative PCR for Ankrd42 **(D)**, Ube3C **(E)**, Slc6a6 **(F)** and Pacrgl **(G)** skiptic exon in spinal cord of Thy1-hTDP-43 mice treated (SAHA; n=4) or not (Vehicle; n=6) with SAHA at disease onset. Data (mean ± SD) are expressed as PEE. **(H-K)** Semi quantitative PCR for Ankrd42 **(H)**, Ube3C **(I)**, Slc6a6 **(J)** and Pacrgl **(K)** skiptic exon in spinal cord of Thy1-hTDP-43 mice treated (SAHA; n=6) or not (Vehicle; n=6) with SAHA at symptomatic stage. Data (mean ± SD) are expressed as PEE. **(L-P)** Semi quantitative PCR for Plod1 **(L)**, Pex16 **(M)**, Ankrd42 **(N)**, Pacrgl **(O)** and Herk2 **(P)** skiptic exon in spinal cord of Thy1-hTDP-43 mice treated (SAHA+Arim.; n=5) or not (Vehicle; n=5) with SAHA+Arim. at symptomatic stage. Data (mean ± SD) are expressed as PEE. (**A-P**) Data are expressed as percentage of Thy1-hTDP-43 mice at PS stage of disease (**A-C**) or vehicle-treated mice (**D-P**). * p ≤ 0.05 by unpaired t test (**E**)

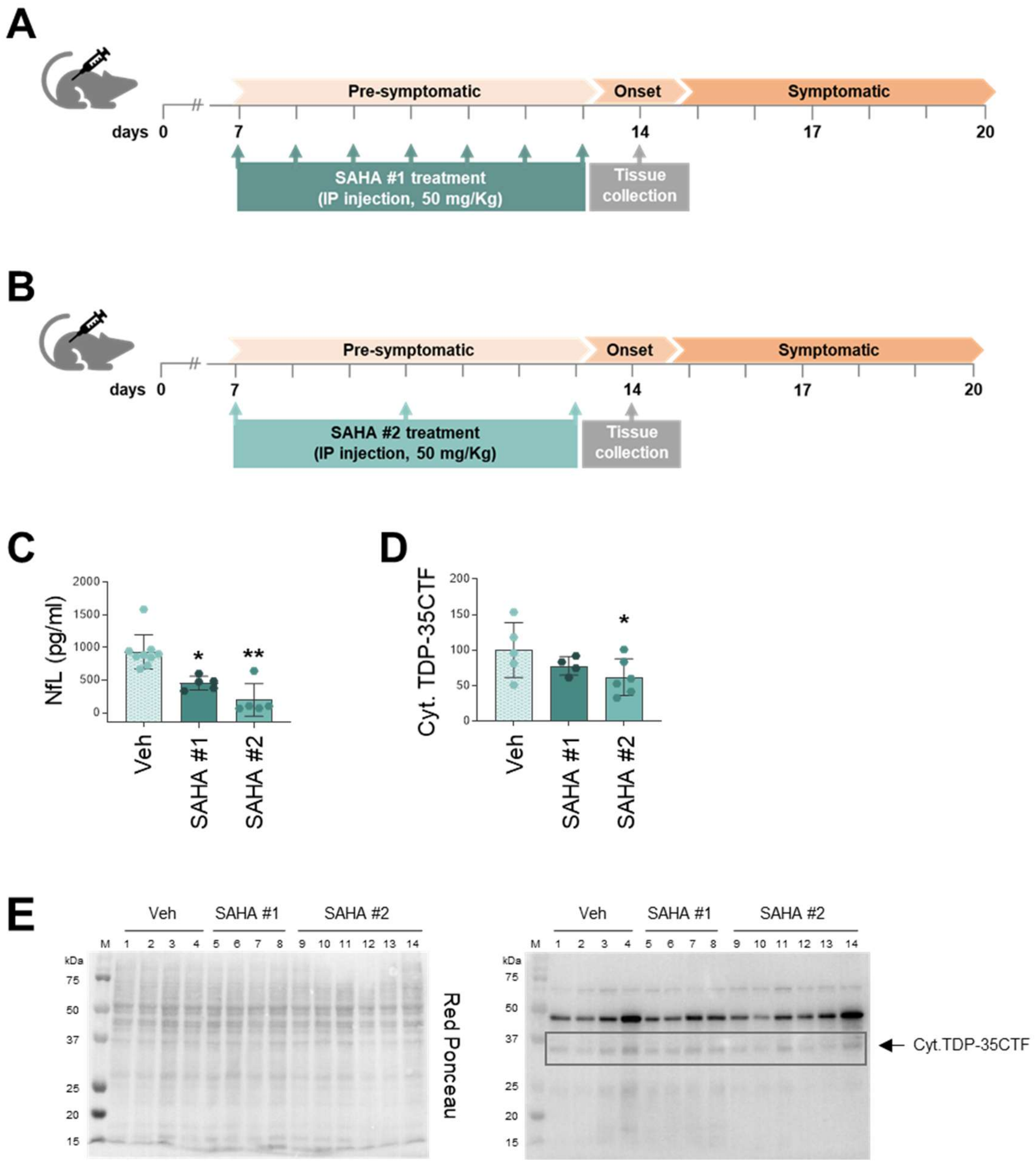


**Supplementary Figure 6. Pilot study to establish optimal treatment schedule for SAHA in homozygous Thy1-hTDP-43 mice.** (**A-B**) Scheme of SAHA treatment in homozygous Thy1-hTDP-43 mice. Mice were treated intraperitoneally with 50 mg/kg of SAHA, from 7 days of age, daily (SAHA #1) (**A**) or every three days (SAHA #2) (**B**) and sacrificed at 14 days for further analysis. (**C**) NfL plasma levels in homozygous Thy1-hTDP-43 mice treated with SAHA daily (SAHA #1; n=5), every three days (SAHA #2; n=5) or vehicle (Veh; n=9). Data are mean ± SD. * $p < 0.05$, ** $p < 0.01$ versus Veh by Kruskal-Wallis with Dunn's multiple comparisons test. (**D**) Levels of cytoplasmic (Cyt.) C-terminal fragment at 35 kDa (TDP-35CTF) in spinal cord of homozygous Thy1-hTDP-43 mice treated with SAHA daily (SAHA #1; n=4), every three days (SAHA #2; n=6) or vehicle (Veh; n=5). Data (mean ± SD) indicates WB immunoreactivity normalized to total protein loading and expressed as percentage of vehicle-treated mice. * $p \leq 0.05$ versus Veh by npaired t test. (**E**) WB after Red Ponceau staining (left) and antibody hybridization (right) for TDP-43, in cytoplasmic fractions derived from spinal cord of Thy1-hTDP-43 treated with SAHA daily (SAHA #1), every three days (SAHA #2) or not treated (Veh). This blot corresponds to the quantitative graphs shown in Figure 4F and Supplementary Figure 6D. Arrow and box indicate the band used for quantification.

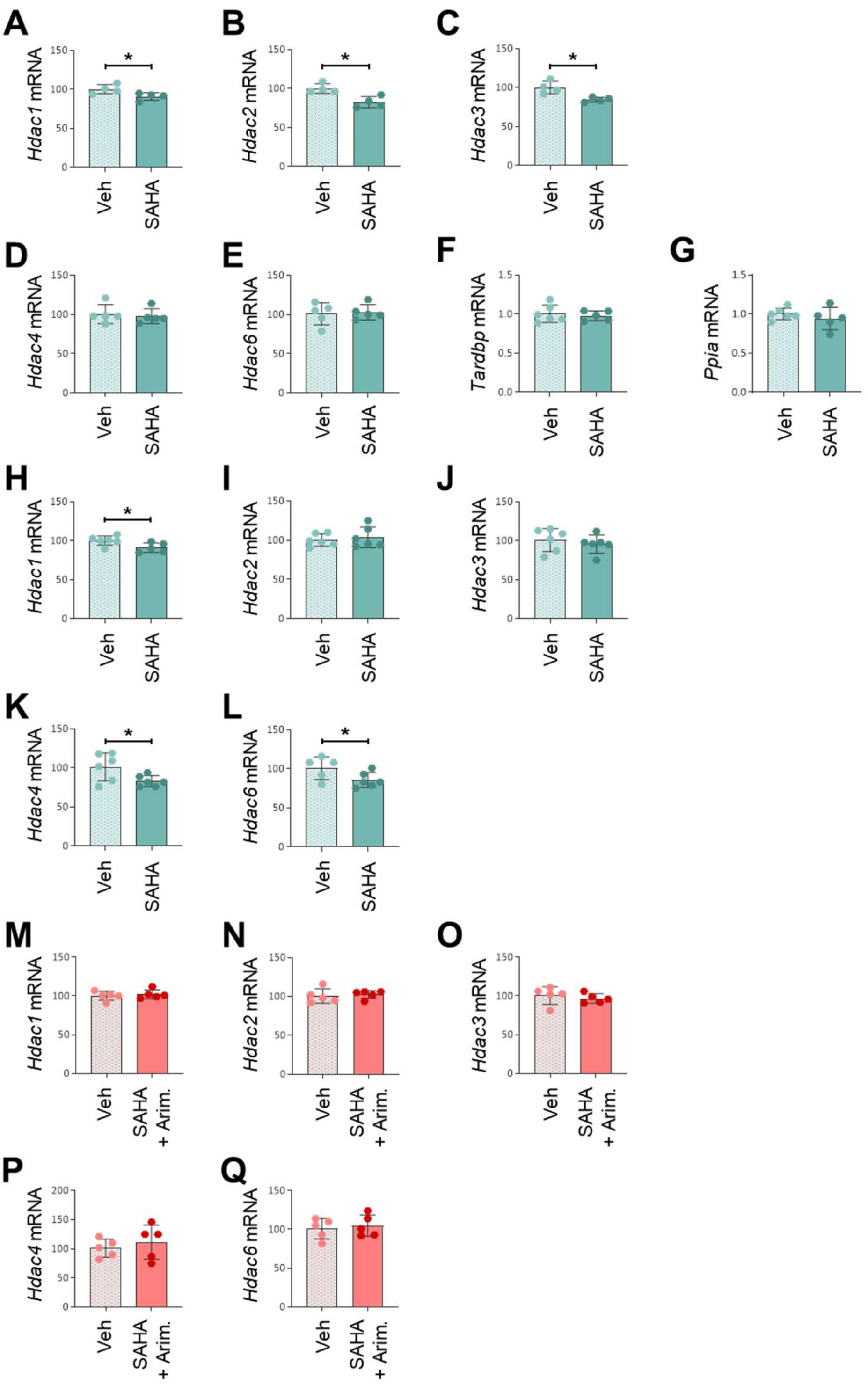


**Supplementary Figure 7. Effect of SAHA and SAHA+Arimoclomol combination on the expression of *Hdacs* in lumbar spinal cord of Thy1-hTDP-43 mice**. (**A-Q**) RT-PCR for *Hdac1*, *Hdac2*, *Hdac3*, *Hdac4*, *Hdac6, Tardbp* and *Ppia* in spinal cord of Thy1-hTDP-43 mice treated with SAHA or vehicle (Veh) and analysed at disease onset (n=4-5 in each experimental group) (**A-G**), at symptomatic stage (n=5-6 in each experimental group) (**H-L**) or treated with SAHA+Arim. combination and analysed at symptomatic stage (n=5 in each experimental group) (**M-Q**). (**A-Q**) Data (mean ± SD) are normalized to *β-actin* and reported as percentage of vehicle-treated mice relative mRNA expression. *p < 0.05 by unpaired t test (**A-C,H,K,L**).

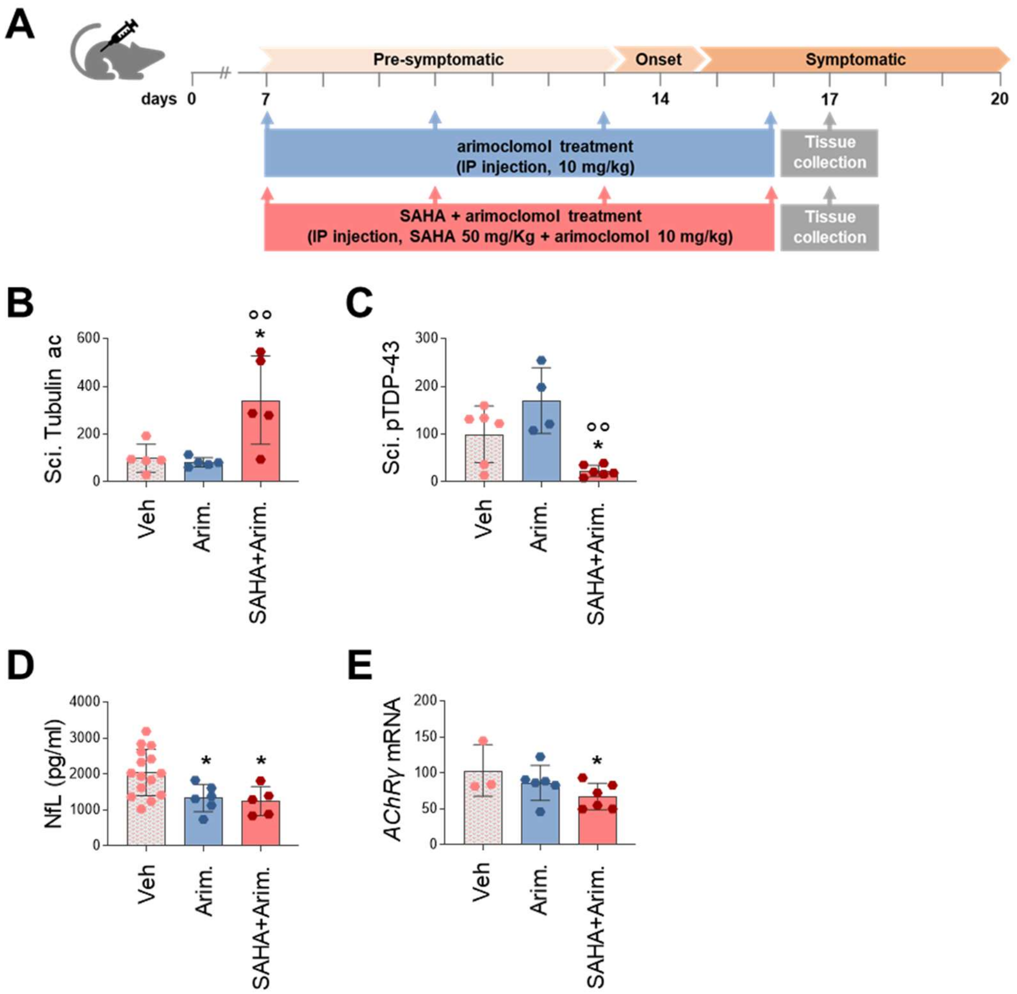


**Supplementary Figure 8. Arimoclomol has no effect on insoluble TDP-43 and markers of neuromuscular pathology. (A)** Scheme of arimoclomol (Arim.) and SAHA+arimoclomol (SAHA+Arim.) combination treatment in homozygous Thy1-hTDP-43 mice. Mice were treated intraperitoneally with 10 mg/kg of Arimoclomol or 50 mg/kg of SAHA and 10 mg/kg of arimoclomol, from 7 days of age, every three days and sacrificed at 17 days for further analysis. **(B-C)** Analysis of acetyl tubulin (Tubulin ac) **(B)** and pTDP-43 **(C)** in sciatic nerve (Sci.) of Thy1-hTDP-43 mice treated with Arim. (n=4-5 mice) or SAHA+Arim (n=6-5 mice) or vehicle (Veh; n=6-5 mice). Data indicates dot blot immunoreactivity normalized to total protein loading **(B-C)** and to tubulin levels **(B)**. *p < 0.05 versus Veh and °°p < 0.01 versus Arim. by one-way ANOVA with Tukey's multiple comparisons. **(D)** NfL plasma levels in Thy1-hTDP-43 mice treated with Arim. (n=6) or SAHA+Arim (n=5) or vehicle (n=15). *p < 0.05 versus vehicle by one-way ANOVA with Tukey's multiple comparisons. **(E)** RT-PCR for *AChR* $\gamma$-subunit in gastrocnemius muscle of Thy1-hTDP-43 mice treated with Arim. (n=6 mice) or SAHA+Arim. (n=6) or vehicle (n=3). Data are normalized to *β-actin* and expressed as relative mRNA expression. *p < 0.05 versus vehicle by unpaired t test. (**B,C,E**) Data are expressed as percentage of vehicle-treated mice.

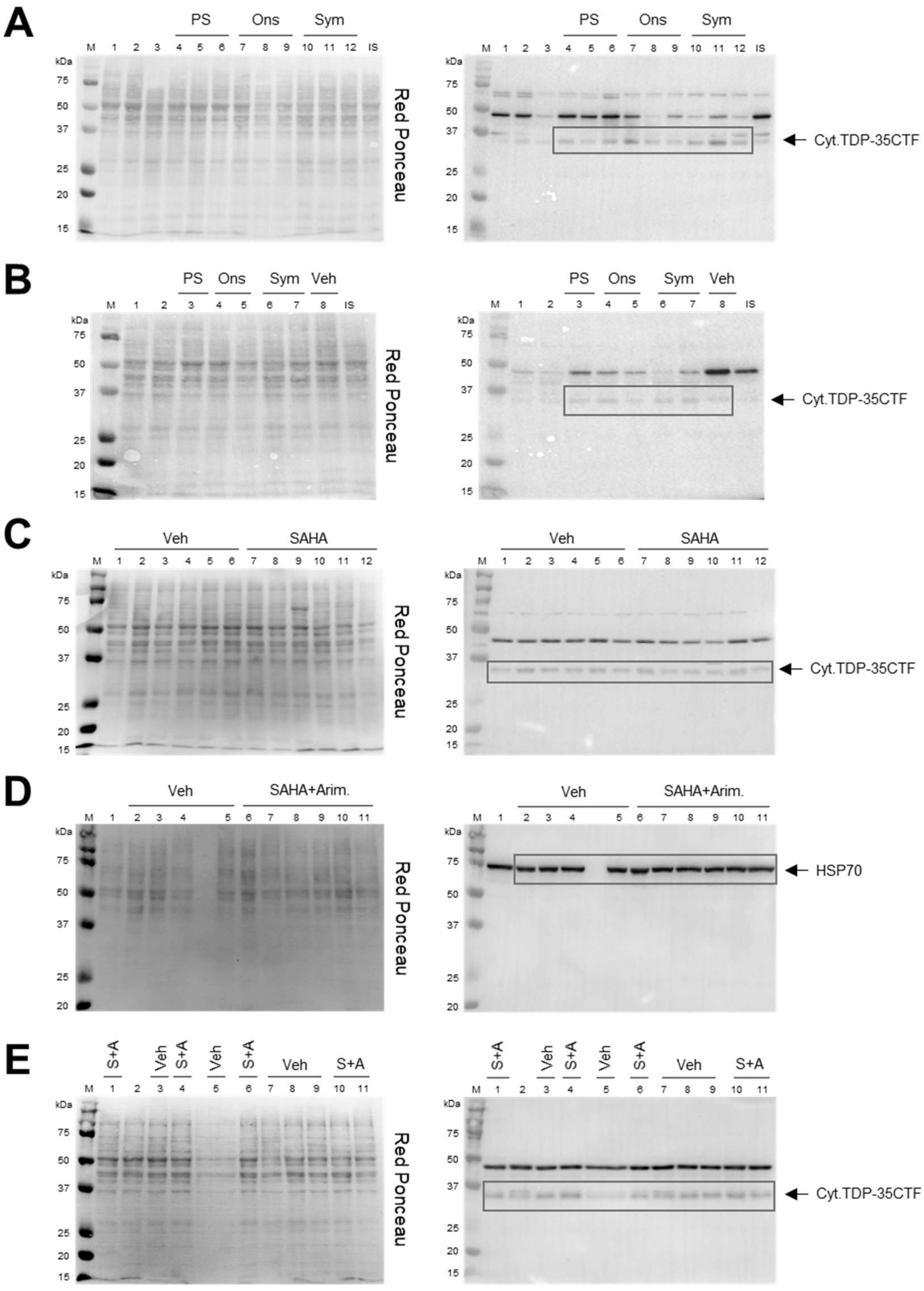


**Supplementary Figure 9. Western blot images supporting Cyt.TDP-43 and HSP70 quantification. (A-B)** WB after Red Ponceau staining (left) and antibody hybridization (right) for TDP-43, in cytoplasmic fraction derived from spinal cord of Thy1-hTDP-43 mice at pre-symptomatic (PS, 7 days of age), onset (Ons, 14 days of age) and symptomatic (Sym, 17 days of age) stage. These blots correspond to the quantitative graph shown in Figure 3E. **(C)** WB after Red Ponceau staining (left) and antibody hybridization (right) for TDP-43, in cytoplasmic fraction derived from spinal cord of Thy1-hTDP-43 treated or not (Veh) with SAHA up to the symptomatic stage. This blot corresponds to the quantitative graph shown in Figure 5G. **(D-E)** WB after Red Ponceau staining (left) and antibody hybridization (right) for HSP70 (**D**) in spinal cord lysate and TDP-43 (**E**) in cytoplasmic fraction derived from spinal cord of Thy1-hTDP-43 treated or not (Veh) with SAHA+arimoclomol (SAHA+Arim.). These blots correspond to the quantitative graphs shown in Figure 6B (D) and Figure 6G (E). **(A-E)** Arrows and boxes indicate the bands used for quantification. IS: internal standard.

**Supplementary Table 1. Skiptic and cryptic primer sequences**

| Gene | Sequence 5'-3' |
|---|---|
| **Ankrd42**[a] | Forward: TCAATGGAGCCAACCTAGCA<br>Reverse: GGCTTGTGTTGCACTGTTCAT |
| **Herc2**[a] | Forward: GGCTGCTAGACCACTCTGAT<br>Reverse: TGACTTTGCCCACATCACCT |
| **Pacrgl**[a] | Forward: GGTCAAGGGTGCTCCTGAAA<br>Reverse: AACCCCTACAGCACTCAACG |
| **Pex16**[a] | Forward: GTGCTGCTCAATGACGGGAT<br>Reverse: ATGCCAGCCTTGAACCAGAT |
| **Plod1**[a] | Forward: CACCTGCTTTCCCTGGATAACT<br>Reverse: ACTGGCCATAGTGTTCCATCTC |
| **Slc6a6**[a] | Forward: CATGACCTCACTGGGAAGCTAT<br>Reverse: ATAGACCAAAAGGTGGGCAGC |
| **Ube3c**[a] | Forward: TGGATGGATCAGAGAGACTGACA<br>Reverse: TGGTTTTAGGACATTCTCCAGC |
| **2610507B11Rik**[b] | Forward: CAGAACACAGAGGTGGAGCAA<br>Reverse: AGAGGTAGCCACCTTAAGAACC |
| **A230046K03Rik**[b] | Forward: CCTTCCTACAGCAGAAAGCTCA<br>Reverse: TGGTGAGGTCTTCAGCAAATGT |
| **Adipor2**[b] | Forward: GCTCAGAAAAGGGCACCAAC<br>Reverse: CGTTCCATAGCATGATGGGC |
| **Adnp2**[b] | Forward: TCCAAAATGTTTCAAATTCCTGTGC<br>Reverse: GGAGAAACATCTCCCCACGA |
| **Celf5**[b] | Forward: CGGAAGCTGTTCGTGGGTAT<br>Reverse: GCCTCTGTGTGGGAAGAAAAC |

[a] skiptic primer sequences

[b] cryptic primer sequences
